# L-Arginine supplementation modulates L-Arg/NO metabolic processes and AMPK/ACC-1 signalling in BNL CL2 hepatocytes

**DOI:** 10.64898/2026.02.03.703662

**Authors:** Saranya Prashath, C Mark Smales

**Affiliations:** School of Natural Sciences, University of Kent, CT2 7NJ, Canterbury, Kent, UK; National Institute for Bioprocessing Research and Training, Co. Dublin, A94 X099, Ireland

**Keywords:** BNL CL2, L-Arg, Nitric Oxide Synthase, Nitric Oxide, AMPK, ACC-1

## Abstract

The enzyme nitric oxide synthase (NOS) breaks down the semi-essential amino acid L-arginine (L-Arg) in the cell to produce citrulline and nitric oxide (NO). NO is a crucial signalling molecule in cells that controls the metabolism of fats and carbohydrates. The aim of this study was to investigate two important genes in the L-Arg-NOS-NO signalling pathway, *AMPK* and *ACC-1*, as markers of the molecular mechanisms that are triggered when liver cells sense elevated L-Arg. Mouse liver epithelial insulin-sensitive BNL CL2 cells were used as a model system and cultured with 0, 400 or 800 µM L-Arg. Cell growth parameters were analysed alongside qRT-PCR based analysis of target transcripts involved in lipid and glucose metabolic pathways. In a further experiment, NOS inhibitor; L-NAME (40 mM) and external NO donor; SNAP (100 µM) were added and the effect on target gene expression analysed. L-Arg addition impacted culture viability and cell growth. AMP-activated protein kinase (AMPK) was regulated in response to L-Arg addition with increasing extracellular concentrations elevating AMPK mRNA and protein expressions. L-NAME decreased target gene expression in an L-Arg addition dependent manner. SNAP (100 µM) addition increased target gene expression after 6 and 24 h. NO, produced as a result of L-Arg addition and the factors L-NAME and SNAP, that regulate NO bioavailability, impacted BNL CL2 cell NO/AMPK/ACC-1 signalling pathways via regulating mRNA expression and subsequently protein expression.

## Introduction

In addition to being the essential building blocks of proteins and peptides, the 20 amino acids are involved in various cellular metabolic processes [1]. The study here focuses on eukaryotic cells, specifically mammalian cells (mouse liver cells), and amino acid metabolism involving the amino acid L-Arg. L-Arg metabolism in the cell can give rise to urea production and ornithine via arginase activity or nitric oxide and L-citrulline production via nitric oxide synthase (NOS) [2].

Calcium-dependent and membrane-associated endothelial NOS (eNOS or NOS3), calcium-dependent neuronal NOS (nNOS or NOS1) [3], and calcium-independent cytosolic inducible NOS (iNOS or NOS2) are the three known isoforms of nitric oxide synthase (NOS) [3–5]. The degree of Ca^2+^ interaction with calmodulin regulates the constitutive low levels of eNOS and nNOS expression in a range of cell types and tissues [5,6]. Under “normal” conditions, iNOS is either not expressed or is expressed at a very low level in cells and tissues [5]. Instead, it’s expression is induced under the appropriate conditions [3]. The three isoforms of NOS are expressed in various tissues, including insulin-sensitive tissues (liver, muscle and adipose tissue) that play an important role in whole-body homeostasis of energy substrates [7].

Tetrahydrobiopterin [(6R)-5,6,7,8-tetrahydrobiopterin] (BH_4_), flavin adenine dinucleotide (FAD), flavin mononucleotide (FMN) and nicotinamide adenine dinucleotide phosphate (NADPH) are co-factors needed to synthesize nitric oxide (NO), a free radical, from L-Arg in the presence of any isoform of NOS [8]. NO is modulated by a range of NOS inhibitors such as N^G^-nitro-L-Arg methylester (L-NAME), N^G^-monomethyl-L-Arg (L-NMMA) and N^G^-nitro-L-Arg (L-NNA) [9].

Nitric oxide is an important intra-and inter-cellular [10,11] signalling molecule that regulates nutrient metabolism [12]. In insulin-sensitive tissues, physiological levels of NO (25–35 μmol) [13] promote glucose uptake and oxidation as well as fatty acid oxidation; in target tissues (such as the liver and adipose), NO inhibits the synthesis of glucose, glycogen and fat and in adipocytes, promotes lipolysis [14]. The production of NO occurs in nearly all cells and tissues of mammals, particularly adipocytes, endothelial cells, cardiac cells, neurons, hepatocytes, myotubes and phagocytic cell [15,16]. The production of nitrites and nitrates, which are oxidative metabolite products of NO, quickly deactivates NO and diffuses into the bloodstream. Because nitrite and nitrate concentrations are commonly employed as markers of NO concentration, they can be used to indirectly estimate the amount of NO that was previously in circulation [9].

The metabolic consequences of excess L-Arg on the NOS/NO signalling pathway in different tissues is not fully mapped. This is somewhat surprising as L-Arg is taken as a dietary supplement to help control body mass/fat matter [17]. Further, the importance of the L-Arg/NOS/NO pathway in insulin sensitizing metabolic pathways has been established *in-vivo* using diabetic and obese animal models. Therefore, in this study, an *in-vitro* cell culture model; BNL CL2 (mouse hepatocytes) has been used to investigate the role of L-Arg/NOS/NO in the regulation of glucose and fatty acid metabolism and signalling in liver cells (BNL CL2) by mapping the responses of key genes and proteins in these pathways that are involved in L-Arg and NO induced signalling cascades.

Furthermore, novel strategies for controlling and preventing diabetes in obese people need be studied and investigated based on less side effects and cost effective. In this study, we aimed to investigate the direct effect of excess L-Arg supplement in insulin sensitive cells (mouse hepatocytes cells) by modulating the expression of key target genes and proteins which are involved in the metabolism of energy substrates in an NO dependent manner. We specifically looked at how excess L-Arg regulates energy metabolism by increasing NO generation, by activating the major energy sensing and downstream genes AMPK and ACC-1, via the AMPK-ACC-1/NOS/NO pathway.

## Results

### Impact of L-Arg addition on culture viability and viable cell numbers in BNL CL2 cells

Samples were collected at 24, 48, 72 and 120 h time points for cell growth analysis of BNL CL2 cells. All cell samples were analysed with biological triplicate cultures. The viable number of cells was compared between the control complete DMEM medium and the no L-arginine SILAC DMEM media (0 µM L-arginine) and when SILAC DMEM had 400 and 800 µM L-arginine added. **Figure 1a** shows that there was a decline in BNL CL2 cell numbers from 24 to 48 h in all samples and conditions. After this time, the proliferation of cells increased for the control and the 400 and 800 µM L-arginine supplemented cell samples from 48 to 72 h. However, the cell number continued to decrease in the no L-Arg (0 µM) cell samples across this time period. Cell number declined in the 400 and 800 µM L-Arg supplemented cell samples between 72 and 120 h, whilst there was a plateau in the growth of the cell samples grown in control complete DMEM. Overall, the viable cell number of BNL CL2 cultures was most impacted by an absence of exogenous L-Arg with minor changes observed between the different concentrations of L-arginine compared to the control. In spite of the similar growth profiles (**Figure 1a**), the effect of L-arginine (0, 400 and 800 µM) compared to the control complete DMEM media was significant (**Figure 1a**) as determined by two-way ANOVA analysis of the means values of the viable cell numbers at the different time points followed by a Tukey multiple comparison test.

**Figure 1.**
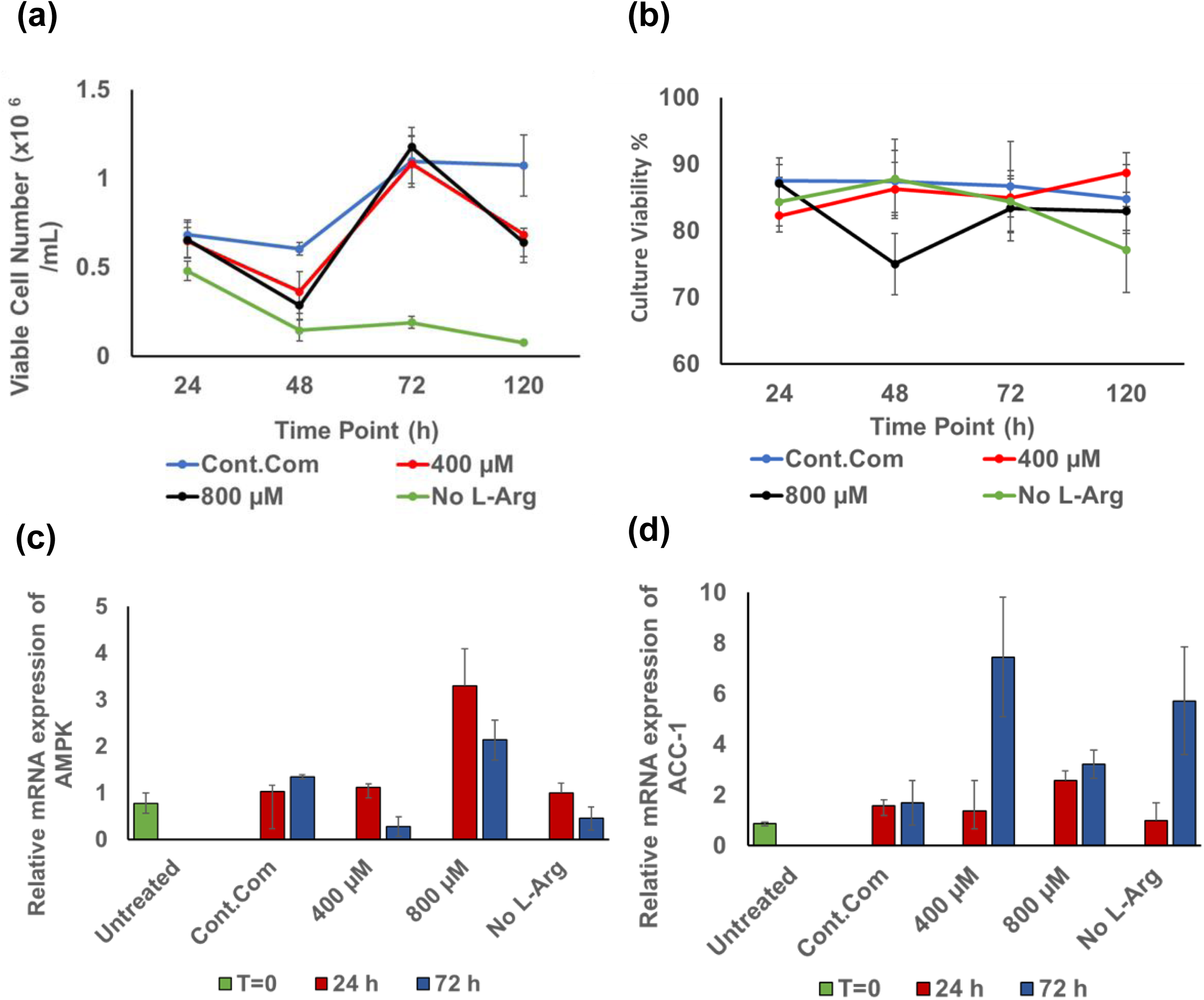
Cell growth and culture viability profiles and relative mRNA expression in BNL CL2 cells with exogenous L-Arg. Cell growth/viable cell number (**a**) and culture viability (**b**) profiles of BNL CL2 cells with different concentrations of L-Arg (0, 400 and 800 µM) and the control complete DMEM media at 24, 48, 72 and 120 h. Relative mRNA expression of AMPK (**c**) and ACC-1 (**d**) cultured in different concentrations of L-Arg in BNL CL2 cells at 24 and 72 h. Data points represent the mean ± SD of each culture sample. Error bars represent the standard deviation from the mean (n = 3).

Figure 1b reports the culture viability of cultures grown in either control complete DMEM or L-arginine supplemented samples (400 and 800 µM) and no L-arginine SILAC DMEM at 24, 48, 72 and 120 h. Culture viability was generally maintained between 80-90% across the 120 h. Noticeably the largest difference was a drop in viability of the cultures with 800 µM L-Arg addition from 24 to 48 h time period, after which the viability then increased again from 48 to 72 h and then remained steady. Overall, the viability of the BNL CL2 cultures was not significantly impacted by the different L-Arg conditions.

### qRT-PCR analysis of AMPK and ACC-1 gene expression in response to L-Arg supplementation of BNL CL2 cell growth media

Relative gene expression analysis (ΔΔCt) was used to quantify target gene mRNA expression. The target gene expression was normalized to β-actin to obtain the ΔCt and then the relative difference (ΔCt) between target and β-actin was normalized to the ΔCt value of the control no L-Arg sample at the 24 h culture time point to obtain the ΔΔCt value. The relative expression of each gene was determined and denoted as RE. The target transcripts AMPK and ACC-1, which are hypothetically modulated by changes in L-Arg through signalling pathways, were investigated.

The first transcript investigated was of the key regulator gene involved in the oxidation of energy substrates, AMPK [18]. AMPK was regulated in response to L-arginine addition with increasing extracellular concentrations increasing the observed AMPK mRNA expression in BNL CL2 cells (Figure 1c). From qRT-PCR data, AMPK gene expression was increased (P <0.0001) in 800 µM L-Arg addition cultures at 24 (RE 3.29) and 72 h (RE 2.14) when compared to the control complete DMEM media cultures. The mRNA levels of the key lipogenic enzyme, ACC1, was decreased (P<0.0001) in cultures with arginine at 400 µM (RE 1.4) compared to the control (RE 1.6) after 24 h but increased (P < 0.0001) at 72 h (RE 7.46) compared to the control complete DMEM media cultures (RE 1.69) (Figure 1d).

### qRT-PCR analysis of AMPK and ACC-1 genes upon L-NAME (4 mM) and SNAP (100 µM) addition to BNL CL2 cell media

The impact of L-NAME (NOS inhibitor) on AMPK gene expression in the cultured BNL CL2 cells with different L-Arg additions with or without L-NAME (4 mM) is presented in Figure 2a. The expression of AMPK mRNA was high (P<0.0001) in samples cultured in no L-Arg with L-NAME at 72 h (2.5-fold compared to the control with L-NAME at 72 h) among all samples with or without L-NAME. It was observed that when cells were treated with 800 µM L-Arg, AMPK gene expression was highest at 24 h (RE 3.3) (Figure 1c). However, when cells were cultured in 800 µM L-Arg with L-NAME, the AMPK expression was decreased (P<0.0001) at 24 h (RE 2.4) in the presence of NOS inhibitor L-NAME, but increased (P<0.0001) in 800 µM L-Arg with L-NAME at 72 h (RE 2.35) compared to arginine at 800 µM at 72 h (RE 2.14).

**Figure 2.**
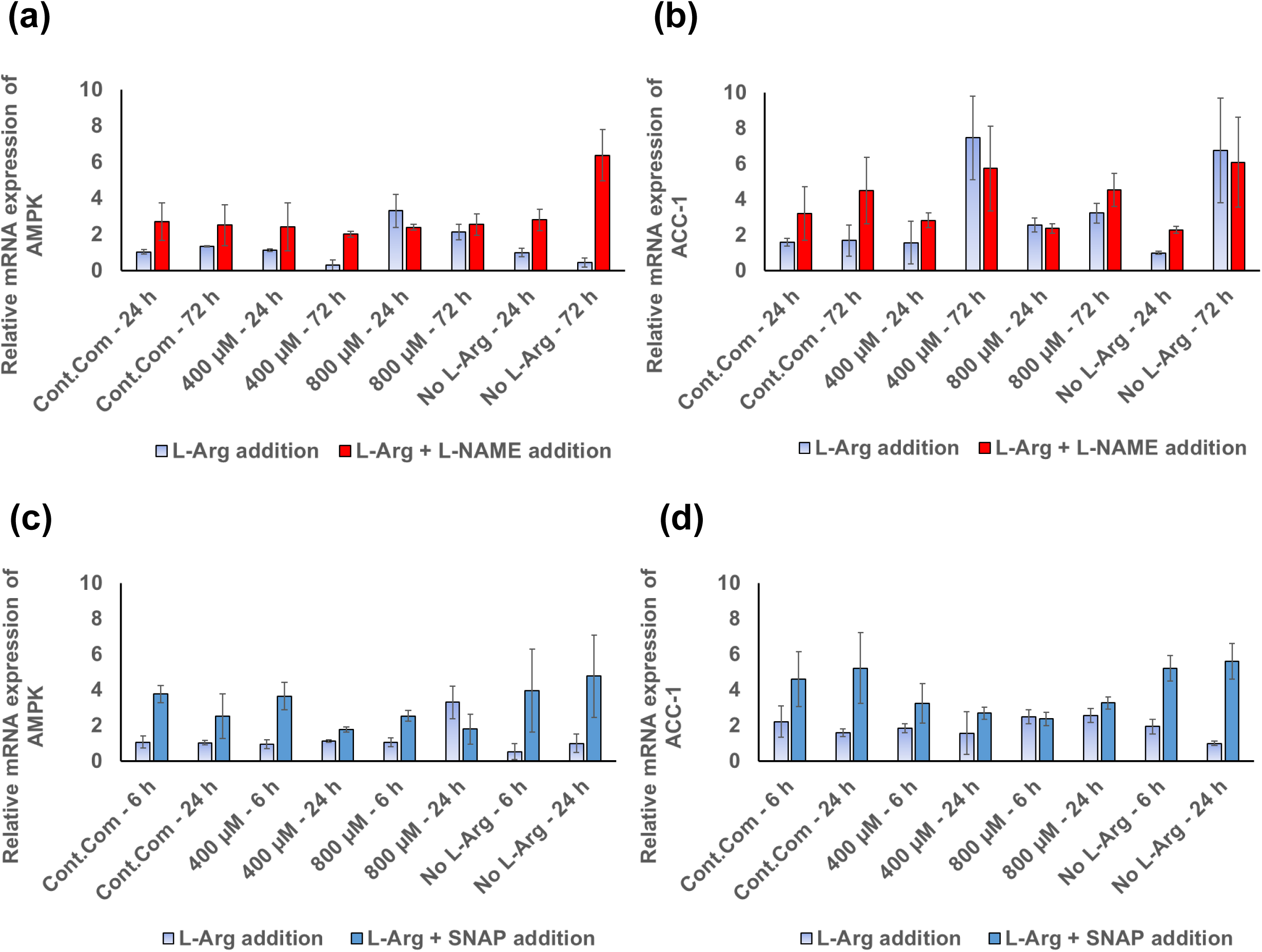
Regulation of relative mRNA expression of AMPK and ACC-1 in BNL CL2 cells with exogenous L-Arg and the modulators L-NAME and SNAP. Relative mRNA transcript expression (ΔΔCt) of AMPK **(a)** and ACC-1 **(b)** in the BNL CL2 cells with nitric oxide synthase inhibitor; L-NAME (4 mM) 24 and 72 h after addition and relative mRNA transcript expression of AMPK **(c)** and ACC-1 **(d)** in the BNL CL2 cells with nitric oxide donor; SNAP (100 µM) 6 and 24 h after addition. All these analyses were undertaken in BNL CL2 cells cultured in medium of different L-Arg concentrations (0, 400 and 800 µM) and control complete DMEM media. Data points represent the mean ± SD of each sample. Error bars represent the standard deviation from the mean (n =3).

Except for L-Arg at 800 µM with L-NAME at 24 h, all samples with L-NAME showed increased (P<0.0001) AMPK gene expression compared to the control samples and the samples treated with different concentrations of L-arginine (0, 400 and 800 µM). AMPK mRNA expression decreased (P<0.0001) with the time of culture when the cells grown in 400 µM L-Arg and L-NAME.

The impact of L-NAME on ACC-1 gene expression is presented in Figure 2b. When comparing the difference of ACC-1 gene expression between L-Arg treated, and L-Arg and L-NAME treated samples, ACC-1 gene expression was increased (P<0.0001) in L-arginine at 800 µM with L-NAME at 72 h (RE 4.53) compared to the ACC-1 expression in the samples cultured in 800 µM L-Arg (RE 3.23). Interestingly, across the time points ACC-1 expression was increased (P<0.0001) in L-Arg and L-NAME treated samples; this was highest in the samples cultured in 0 µM L-Arg and L-NAME (24h; RE 2.3 and 72 h; RE 6.07). The highest ACC-1 gene expression was in 0 µM L-Arg and L-NAME samples at 72 h (1.35-fold) compared to the control sample cultured in complete DMEM with L-NAME.

AMPK mRNA expression was evaluated in excess exogenous L-arginine with or without SNAP (NO donor; 100 µM) addition and the relative expression is presented in Figure 2c. Overall, the samples cultured in L-Arg and SNAP had increased (P<0.0001) expression of AMPK at 6 and 24 h compared to the samples cultured in L-Arg without SNAP at the same time points. However, AMPK mRNA levels decreased (P<0.0001) at 800 µM with SNAP at 24 h (0.54-fold) in comparison to the 800 µM L-Arg. After 6 h, when compared with and without SNAP addition, there was increased (P<0.0001) AMPK mRNA expression in the samples cultured with NO donor SNAP (the control complete DMEM + SNAP; 4.34-fold, 400 µM + SNAP; 3.84-fold, 800 µM + SNAP; 2.4-fold and 0 µM + SNAP; 7.43-fold). Noticeably, this increase was highest in 0 µM L-Arg (7.43-fold) among all the samples at 6 h.

The ACC-1 mRNA expression profile is presented in Figure 2d. Overall, SNAP addition increased the expression of ACC-1 mRNA in L-Arg treated hepatocytes cells at 6 and 24 h in comparison to the samples cultured in different concentrations of L-Arg and the control complete DMEM. Comparison of ACC-1 mRNA expression in the presence and absence of SNAP in the L-Arg treated BNL CL2 cells showed that there was an increase (P<0.0001) between the samples cultured with SNAP compared to the samples without SNAP at 6 h (the control compete DMEM + SNAP; 2.1-fold, 400 µM + SNAP; 1.76-fold and 0 µM + SNAP; 2.69-fold), except in 800 µM and SNAP (0.95-fold). Samples cultured with SNAP and L-Arg had increased (P<0.0001) ACC-1 mRNA expression with culture time, except at 400 µM L-Arg with SNAP (P<0.0001) (6 h; RE 3.25 and 24 h; RE 2.7).

### AMPK and ACC-1 protein and their post-translational phosphorylation level in response to L-Arg supplementation to BNL CL2 cells

Western blot analysis was used to compare protein expression and phosphorylation of target proteins (Figures 3a**-3h**). Target protein expression was normalized to a reference protein; β-actin. All data was then normalized to the value of no L-Arg added samples (0 µM L-Arg) at 24 h culture time point, denoted as RE in the following sections.

**Figure 3.**
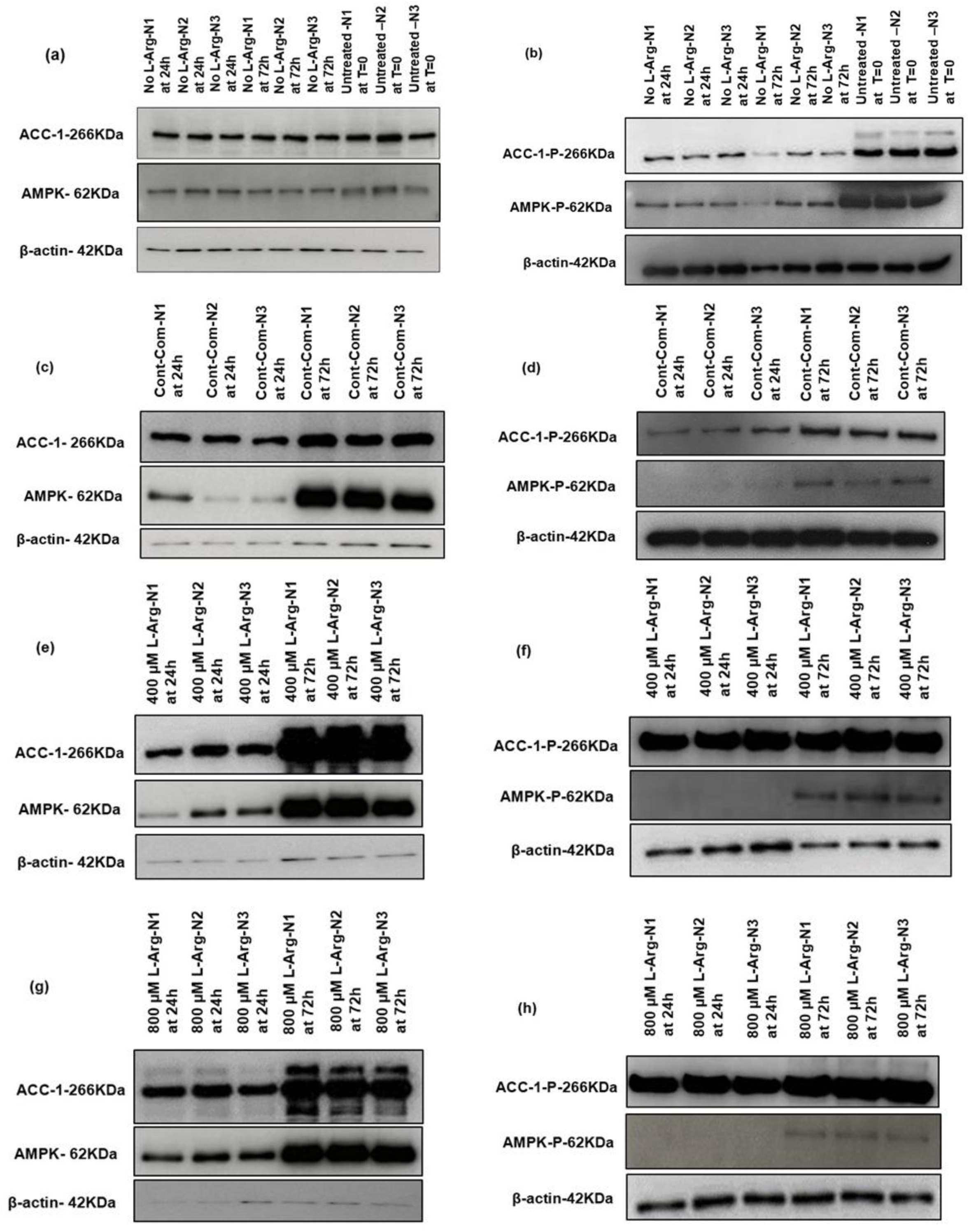
Western blot comparison for target proteins and phosphorylated proteins expression in BNL CL2 cells with exogenous L-Arg. Western blot comparison of the expression of AMPK and ACC-1 proteins **(a, c, e and g)** involved in L-Arg/NO metabolic pathway signalling, and the amount of phospho-protein **(b, d, f and h)** of these targets in BNL CL2 cells cultured in no L-Arg SILAC DMEM media, control complete DMEM media and 400 and 800 µM L-Arg in L-Arg free SILAC DMEM media for 24 and 72 h and in one of the control, untreated culture samples at T=0. β-actin is used as a loading control. The same amount of protein (10 µg) from different treatment groups was loaded from biological triplicate cultures into 10% SDS–polyacrylamide gels for the separation of target proteins.

A comparison across L-Arg concentrations with culture time of total (Figure 4a) and phosphorylated (Figure 4b) levels of AMPK in BNL CL2 cells was analysed. The AMPK protein showed increased expression (P < 0.0001) when cultured in 400 (2.6-fold) and 800 µM (2.75-fold) L-arginine compared to the control complete media samples at 24 h. AMPK protein expression peaked (1.52-fold) in 400 µM cultured samples at 72 h.

**Figure 4.**
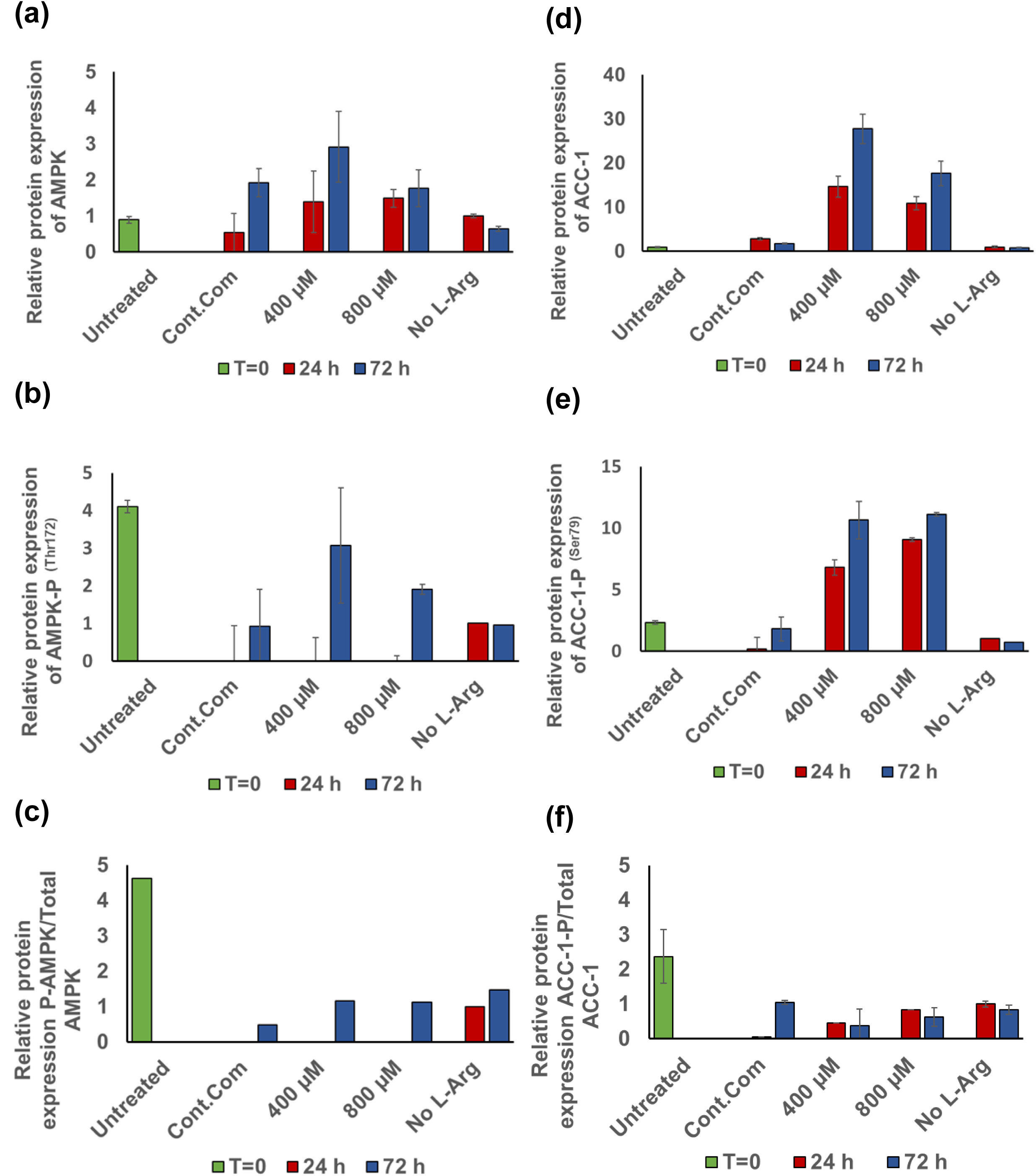
Relative total and phosphorylated protein expressions of AMPK and ACC-1 in BNL CL2 cells with exogenous L-Arg Relative protein amounts for total AMPKα **(a)** and ACC-1 **(b)**, phosphorylated AMPKα at Thr172 (AMPKα-P) **(c)** and ACC-1 at Ser79 (AMPKα-P) **(d)** and the ratio of AMPKα-P to total AMPKα in BNL CL2 cells **(e)** and ACC-1-P to total ACC-1 in BNL CL2 cells **(f)**. Cells were cultured for 24 or 72 h in customized media containing 0, 400 and 800 µM L-Arg. Controls were addition of complete DMEM or untreated cultures at T=0. Bands in the blots were quantified using ImageJ software and the data are normalised to β-actin and then expressed as relative to the no L-Arg SILAC DMEM (0 µM L-Arg) control at 24 h. Data points represent the mean ± SD. Error bars represent the standard deviation from the mean (n = 3).

When phosphorylation of AMPK was investigated Figure 4b, in 400 (0.7-fold) and 800 µM (0.46-fold) L-Arg cultured samples phosphorylation was decreased (P < 0.0001) when compared to untreated samples at T=0 (RE 4.1). There were no differences (P < 0.0001) in phosphorylated AMPK protein in no L-Arg media samples. Increasing L-arginine concentrations from 400 to 800 μM resulted in decreased (P < 0.0001) phosphorylated AMPK levels at 72 h. A summary of the impact of L-arginine concentration on AMPK protein expression and phosphorylation with culture time is reported in Figures 4a **-4c**.

Total and phosphorylated levels of ACC-1 protein in BNL CL2 cells under different L-arginine culture conditions are presented in Figures 4d **-4f**. The expression of this regulator of fatty acid metabolism, ACC-1, increased (P<0.0001) in both total protein and phosphorylated ACC-1 in cells cultured in L-arginine at 400 and 800 μM. Total ACC-1 protein was increased in 400 (5-fold) and 800 µM (4-fold) L-arginine cultures (P<0.0001) compared to the control complete DMEM addition cultures at 24 h. After 72 h of culture time the increase in expression was further elevated in 400 and 800 μM L-arginine cultures (P<0.0001) with total ACC-1 amounts increased by 16-fold and 10-fold, respectively, in comparison with control complete DMEM cultures. Total ACC-1 protein was decreased (P<0.0001) in the untreated sample at T=0 (RE-0.98) and no L-Arg added samples had the lowest relative expression of ACC-1 at 72 h (RE 0.8) among all samples.

When phosphorylation of ACC-1 was investigated (Figure 4e), this was much higher (P<0.0001) in 400 and 800 µM L-arginine cultured cells at 24 and 72 h compared to the control complete media cultures. The ratio of phosphorylated ACC-1 to total ACC-1 is shown in Figure 4f.

### AMPK and ACC-1 protein and their post-translational phosphorylation levels in response to L-Arg supplementation in BNL CL2 cells with L-NAME (4 mM) and SNAP (100 µM)

Protein expression (AMPK and ACC-1) was also investigated upon L-NAME (Figures 5a **-5d**) and SNAP (Figures 6a **-6d**) additions to BNL CL2 cells. Figure 5e presents the relative protein expression of AMPK upon L-NAME addition to BNL CL2 cells in different exogenous L-arginine concentrations. Interestingly, AMPK protein expression was decreased (P<0.0001) in the samples cultured in L-NAME and L-arginine (400 and 800 µM) at 24 (0.26-fold and 0.31-fold, respectively) and 72 h (0.13-fold and 0.24-fold, respectively) compared to the arginine at 400 and 800 µM samples. Noticeably, the relative AMPK protein expression in the samples cultured in L-NAME and excess exogenous L-arginine (400 and 800 µM) was more or less the same irrespective of the time points (for 400 µM + L-NAME at 24; RE 0.36 and 72 h; RE 0.37 and for 800 µM + L-NAME at 24; RE 0.46 and 72 h; RE 0.42).

**Figure 5.**
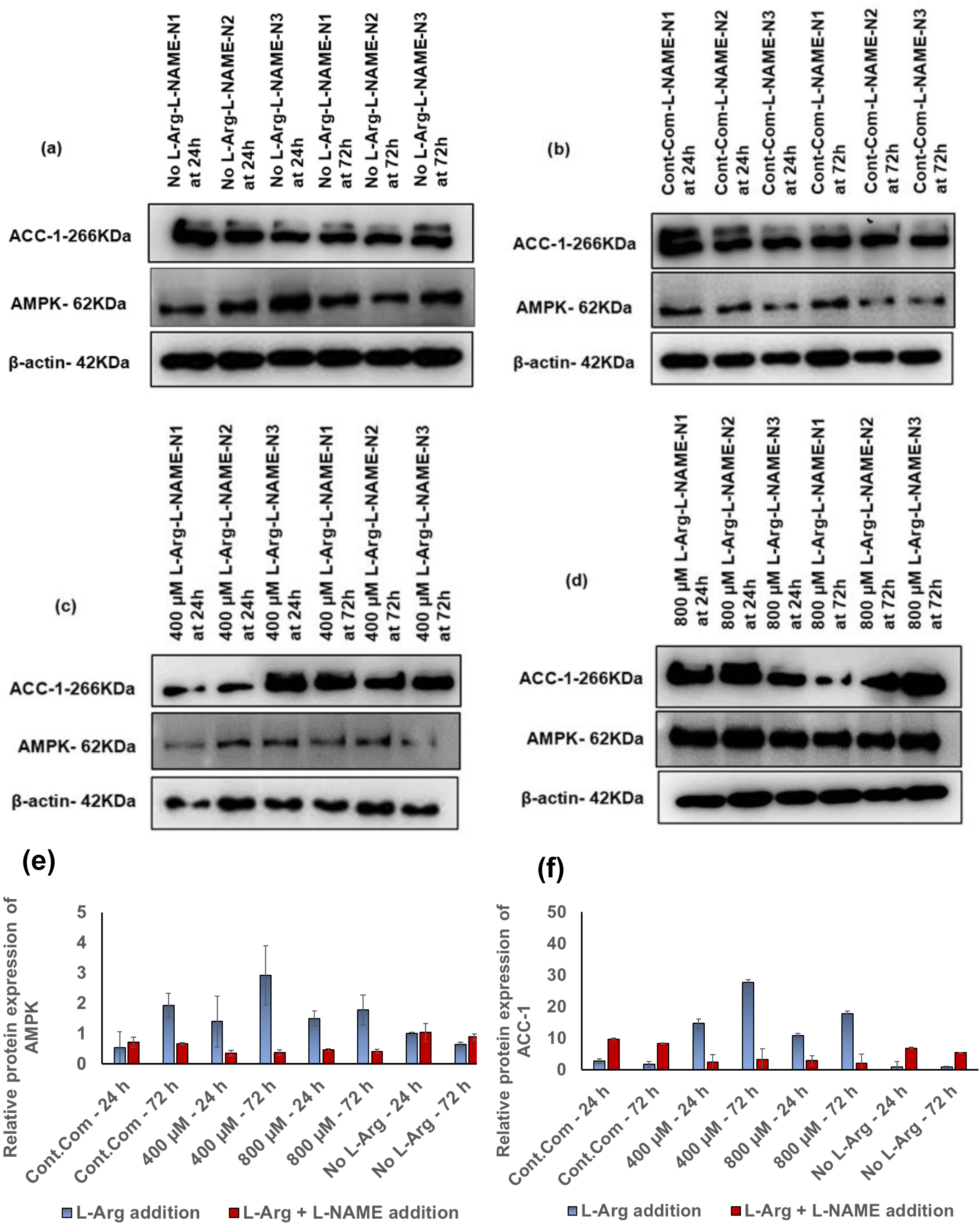
Western blot and relative AMPK and ACC-1 protein expressions in BNL CL2 cells with exogenous L-Arg and L-NAME (4 mM) Western blot comparison of the expression of key protein; AMPK and lipogenic ACC-1 protein levels investigated in L-Arg/NO metabolic pathway signalling in BNL CL2 cells cultured in customized media containing **(a)** no L-Arg SILAC DMEM and L-NAME (4 mM), **(b)** complete DMEM and L-NAME (4 mM), **(c)** 400 µM L-Arg and L-NAME (4 mM), and **(d)** 800 µM L-Arg and L-NAME (4 mM) for 24 and 72 h. Relative protein expression of AMPK **(e)** and ACC-1 **(f)** involved in L-Arg/NO metabolic pathway signalling in BNL CL2 cells cultured in different concentrations of L-Arg and the control with or without L-NAME for 24 and 72 h conditions investigated. β-actin is used as a loading control. The same amount of protein (10 µg) from different treatment groups was loaded from biological triplicate cultures into 10% SDS–polyacrylamide gels for the separation of target proteins. Data points represent the mean ± SD. Error bars represent the standard deviation from the mean (n = 3).

**Figure 6.**
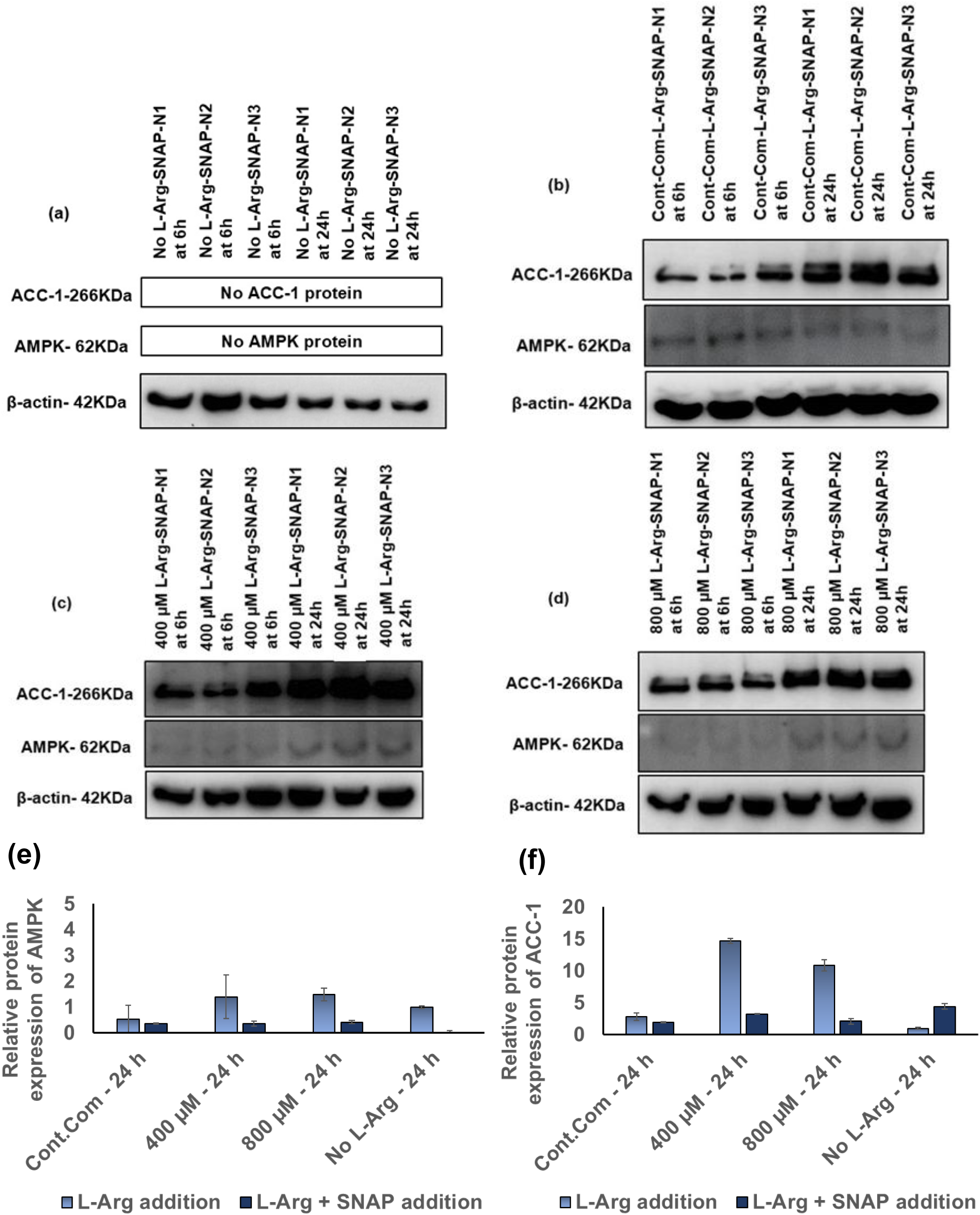
Western blot and relative AMPK and ACC-1 protein expressions in BNL CL2 cells with exogenous L-Arg and SNAP (100 µM) Western blot comparison of the expression of key protein; AMPK and lipogenic ACC-1 protein levels investigated in L-Arg/NO metabolic pathway signalling in BNL CL2 cells cultured in customized media containing **(a)** no L-Arg SILAC DMEM and SNAP (100 µM) **(b)** complete DMEM and SNAP (100 µM), **(c)** 400 µM L-Arg and SNAP (100 µM), and **(d)** 800 µM L-Arg and SNAP (100 µM) for 6 and 24 h. Relative protein expression of AMPK **(e)** and ACC-1 **(f)** involved in L-Arg/NO metabolic pathway signalling in BNL CL2 cells cultured in different concentrations of L-Arg and the control with or without SNAP for 24 h conditions investigated. β-actin is used as a loading control. The same amount of protein (10 µg) from different treatment groups was loaded from biological triplicate cultures into 10% SDS–polyacrylamide gels for the separation of target proteins.

Figure 5f presents the relative protein expression of ACC-1 upon L-NAME addition. Relative ACC-1 protein expression was unexpectedly increased to the highest levels in the samples cultured in L-Arg at 24 (400 µM; 5.1-fold and 800µM; 3.8-fold) and 72 h (400 µM; 16.1-fold and 800µM; 10.3-fold) compared to the control complete DMEM. However, when the NOS inhibitor L-NAME was added to these samples the relative ACC-1 expression was decreased (P<0.0001) to the lowest levels at 24 (400 µM; 0.24-fold and 800µM; 0.3-fold) and 72 h (400 µM; 0.4-fold and 800µM; 0.25-fold) compared to the L-NAME added to the control complete DMEM. Like complete DMEM with L-NAME at 24 and 72 h, the samples cultured in 0 µM L-Arg showed increased (P<0.0001) ACC-1 protein expression at 24 (RE 6.89) and 72 h (RE 5.44) compared to the L-Arg (400 and 800 µM) and L-NAME samples at the same time points. Overall, L-NAME addition to the samples cultured in L-Arg had decreased (P<0.0001) relative ACC-1 protein expression, whereas L-NAME addition to the control complete DMEM and no L-Arg had increased (P<0.0001) relative ACC-1 protein expression.

Relative AMPK protein expression in BNL CL2 cells cultured with exogenous L-Arg and SNAP at 24 h time point is presented in Figure 6e. After 24 h, AMPK protein expression was decreased (P<0.0001) in the samples cultured with SNAP (the control + SNAP 0.41-fold, 400 µM + SNAP; 0.2-fold and 800 µM 0.23-fold) compared to the samples cultured without SNAP. The samples cultured with 0 µM L-Arg and SNAP had the lowest AMPK expression at 24 h.

Relative ACC-1 protein expression in BNL CL2 cells cultured with exogenous L-Arg and SNAP at 24 h time point is presented in Figure 6f. Like AMPK expression, after 24 h, the ACC-1 protein expression was decreased (P<0.0001) in samples cultured with SNAP compared to the samples cultured without SNAP. There was a large decrease (P<0.0001) in ACC-1 protein expression in the samples cultured in 400 and 800 µM with SNAP. The samples cultured with 0 µM L-Arg and SNAP had highest (P<0.0001) ACC-1 expression at 24 h (RE 3.0) among the SNAP treated samples.

### Determination of nitric oxide / nitrite measurements by Griess Assay in BNL CL2 cells grown in different concentrations of exogenous L-Arg

The amount of nitrite present in samples was normalised to the amount of nitrite present in the no L-Arg (0 µM) cell samples at 24 h and then the relative (Figures 7a, 7c and 7e) and the actual (Figures 7b, 7d and 7f) amount of nitrite present in the cell samples are presented Figures 7a**-7f**. Interestingly, increasing extracellular concentrations of L-arginine to 800 µM increased NO synthesis in BNL CL2 cells (Figure 7a **and 7b**). This is consistent with a reported study where cultured primary rat hepatocytes cells were incubated with or without L-arginine medium (L-arginine content 50 mg/L, 287 µM) and increased NO formation via the NO/cyclicGMP pathway was reported in the L-arginine cultured samples [19]. The amount of NO_2_^-^present in 800 µM L-Arg samples was high (P<0.0001) at 24 h (2.5-fold) compared to the control complete DMEM cultured samples at the same time point. It is noted that in regard to these findings, the transcript analysis described earlier in Figure 1c shows that the relative gene expression of AMPK was increased in cells cultured in 800 µM L-arginine at 24 h when the amount of nitrite present in the samples was high.

**Figure 7.**
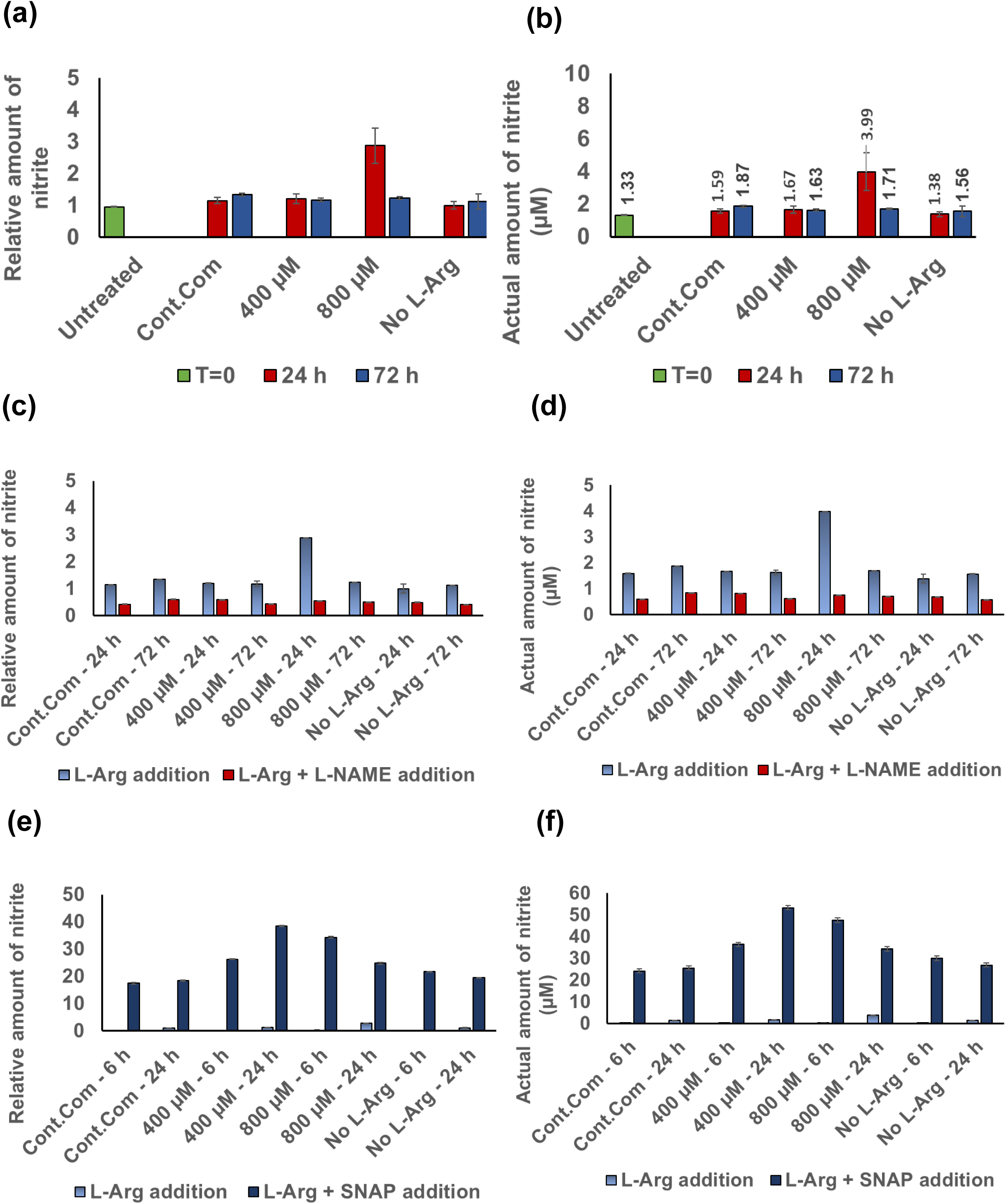
The effect of exogenous L-Arg concentration, NOS inhibitor (L-NAME) and external NO donor (SNAP) on nitrite production in BNL CL2 cells. Cell culture supernatant was obtained from cultured BNL CL2 cells grown in the presence of 0, 400 or 800 µM L-Arg and the controls complete DMEM addition for 24 or 72 h and untreated cell samples at T=0 (relative **(a)** and actual **(b)** values), cells grown in the presence of L-NAME (4 mM, relative **(c)** and actual **(d)** values) and cells grown in the presence of SNAP (100 µM) in 0, 400 or 800 µM L-Arg and the control complete DMEM for 6 or 24 h (relative **(e)** and actual **(f)** values). Quantified nitrite was normalised to the nitrite amount presence in cultures with no L-Arg SILAC DMEM (0 µM L-Arg) at 24 h. Data points represent the mean ± SD. Error bars represent the standard deviation from the mean (n = 3).

### Determination of nitric oxide / nitrite measurements by Griess Assay in BNL CL2 cells grown in different concentrations of exogenous L-Arg with L-NAME (4 mM) and SNAP (100 µM)

When L-NAME (4 mM) was added to the BNL CL2 cells cultured in L-Arg (0, 400 and 800 µM) and the control complete DMEM, the amount of nitrite present is reported in Figures 7c **and 7d**. The amount of signalling molecule, NO involved in L-Arg/NO pathway decreased (P<0.0001) in the samples cultured with L-NAME after 24 (the control + L-NAME; 0.37-fold, 400 µM + L-NAME; 0.49-fold, 800 µM + L-NAME; 0.19-fold and 0 µM + L-NAME; 0.49-fold) and 72 h (the control + L-NAME; 0.44-fold, 400 µM + L-NAME; 0.37-fold, 800 µM + L-NAME; 0.41-fold and 0 µM + L-NAME; 0.36-fold) compared to the samples cultured without L-NAME. Interestingly, a large decrease (P<0.0001) in the amount of nitrite was observed in the samples cultured at highest L-Arg 800 µM with L-NAME (0.54 µM nitrite) in comparison to the samples cultured without L-NAME (2.88 µM nitrite) after 24 h.

When SNAP (100 µM) was added, the amount of nitrite present in the cell samples is shown in Figures 7e **and 7f**. Overall, upon addition of the NO donor to cells cultured in L-Arg (0, 400 and 800 µM) and the control complete DMEM there was a large increase (P<0.0001) in the nitrite amount at 6 (the control + SNAP; 623.51-fold, 400 µM + SNAP; 571.52-fold; 800 µM + SNAP; 180.58-fold and 0 µM + SNAP; 542.25-fold) and 24 h (the control + SNAP; 16.02-fold, 400 µM + SNAP; 32.08-fold; 800 µM + SNAP; 8.60-fold and 0 µM + SNAP; 19.44-fold) compared to the cell samples cultured without NO donor at the same timepoints. The highest amount of nitrite present in the samples was that of the L-arginine at 400 µM with SNAP (53.36 µM) compared to the 400 µM L-Arg without SNAP at 24 h (1.67 µM).

### HPLC analysis of residue L-arginine, L-ornithine and L-citrulline in the cell culture media of BNL CL2 cells cultured in different initial L-Arg concentrations

The amount of each target amino acid present in samples was normalised to the amount of particular amino acid present in the no L-Arg (0 µM) cell samples at 24 h and then the relative and the actual amount of target amino acids present in the cell samples are presented in the Figures 8a**-8f**. When investigating L-arginine present in the cultured serum samples (Figures 8a **and 8d**) the amount of L-arginine increased (P<0.0001) in the samples cultured in control complete DMEM (956.3 µM) and 400 µM L-arginine (981.1 µM) at 72 h time point. Arginine at 400 µM had decreased (0.87-fold, P<0.0001) amount of L-arginine at 24 h but the amount was increased (1.025-fold, P<0.0001) at 72 h in comparison to the control. The opposite profile was observed in samples cultured in 800 µM L-arginine, which was increased (1.017-fold, P<0.0001) in the amount of L-arginine at 24 h after which this was decreased (0.52-fold, P<0.0001) at 72 h compared to the control.

**Figure 8.**
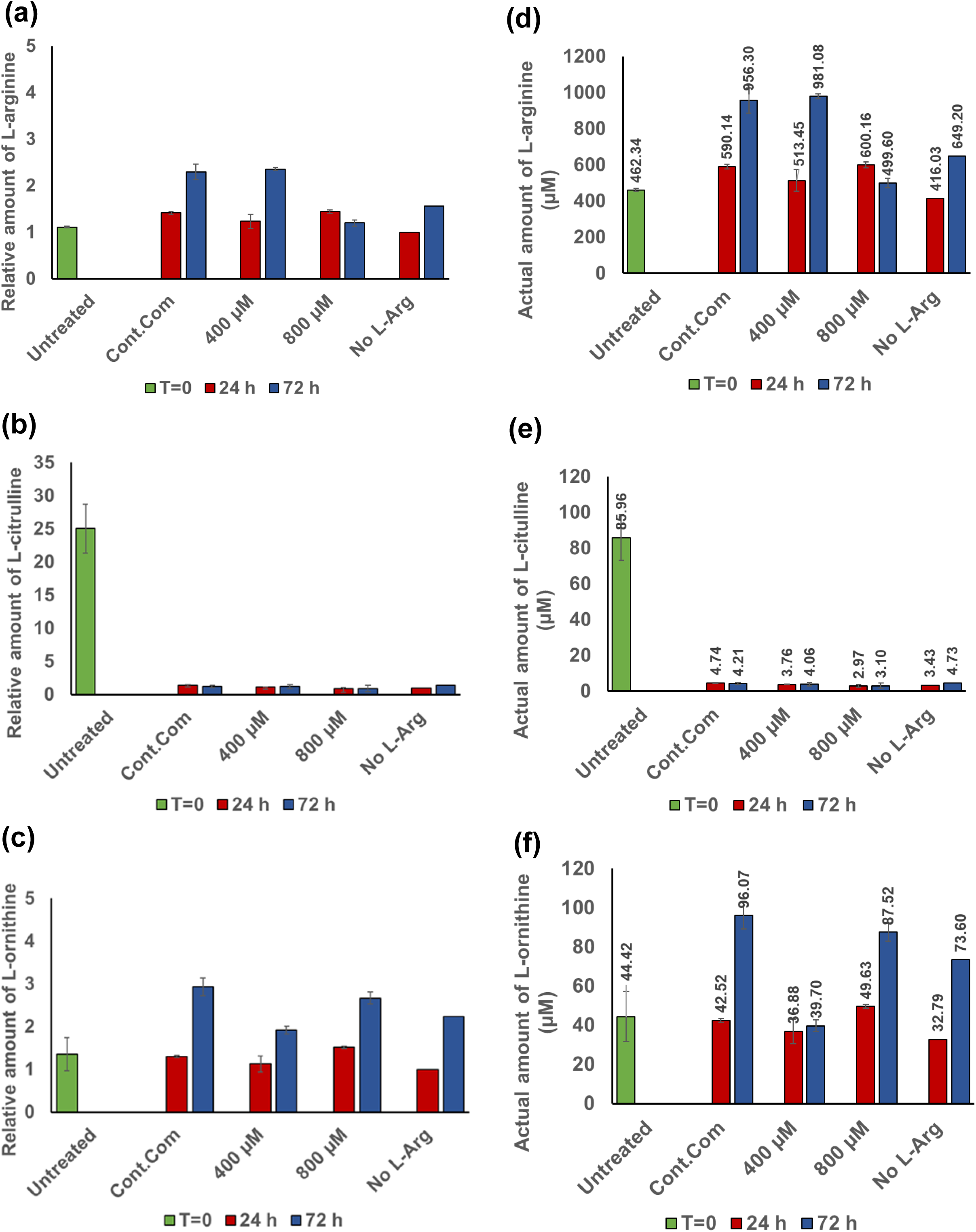
Residual serum L-Arg, L-Cit and L-Orn in BNL CL2 cells with exogenous L-Arg. Relative and actual residue amount of L-Arg **(a and b)**, L-Cit **(c and d)** and **(e and f)** analysed in plasma samples of BNL CL2 cells cultured in different amount of L-Arg (0, 400 and 800 µM) and the control complete DMEM media after 24 and 72 h. Untreated cultures at T=0. Data points represent the mean ± SD of each sample. Error bars represent the standard deviation from the mean (n =3).

When analysing L-citrulline present in the different cultured conditions (Figures 8b **and 8e**), surprisingly, untreated sample at T=0 showed the highest amount of L-citrulline (85.96 µM) (P<0.0001) among all samples. The second highest amount of L-citrulline was in samples cultured in control complete DMEM at 24 h (4.74 µM) and no L-arginine added at 72 h (4.73 µM). More or less the same amount of L-citrulline (P<0.0001) was present in 400 and 800 µM L-arginine samples across culture.

When investigating the amount of L-ornithine present in the cultured samples (Figures 8c **and 8f**), there was an overall increase (P<0.0001) in the amount of L-ornithine across the time points in all samples. Samples cultured in L-arginine at 400 µM contained decreased (P<0.0001) amounts of L-ornithine at 24 (0.87-fold) and 72 h (0.655-fold). However, in samples cultured in L-Arg at 800 µM there was increased (P<0.0001) amounts of L-ornithine present at 24 (1.17-fold) and 72 h (0.91-fold).

### HPLC analysis of residue L-arginine, L-ornithine and L-citrulline in the cell culture media of BNL CL2 cells cultured in different initial L-Arg with L-NAME (4 mM) and SNAP (100 µM)

L-arginine present in the cell culture samples (Figures 9a **and 9d**) collected from cells cultured in 400 and 800 µM L-Arg with L-NAME after 24 (5.37-fold and 4.89-fold, respectively) and 72 h (5.58-fold and 5.98-fold, respectively) was high (P<0.0001) compared to the samples cultured without L-NAME at the same time points. The amino acid synthesised via the L-Arg/NOS metabolic pathway, L-citrulline was analysed in the cell culture supernatant and is shown in (Figures 9b **and 9e**). The amount of L-citrulline was decreased (P<0.01) in the samples cultured in L-Arg (400 and 800 µM) and L-NAME (0.36-fold and 0.47-fold, respectively) compared to the samples without L-NAME at 24 h. After 72 h the amount of L-citrulline decreased (P<0.01) in L-Arg at 400 µM with L-NAME (0.78-fold), over samples cultured without L-NAME.

**Figure 9.**
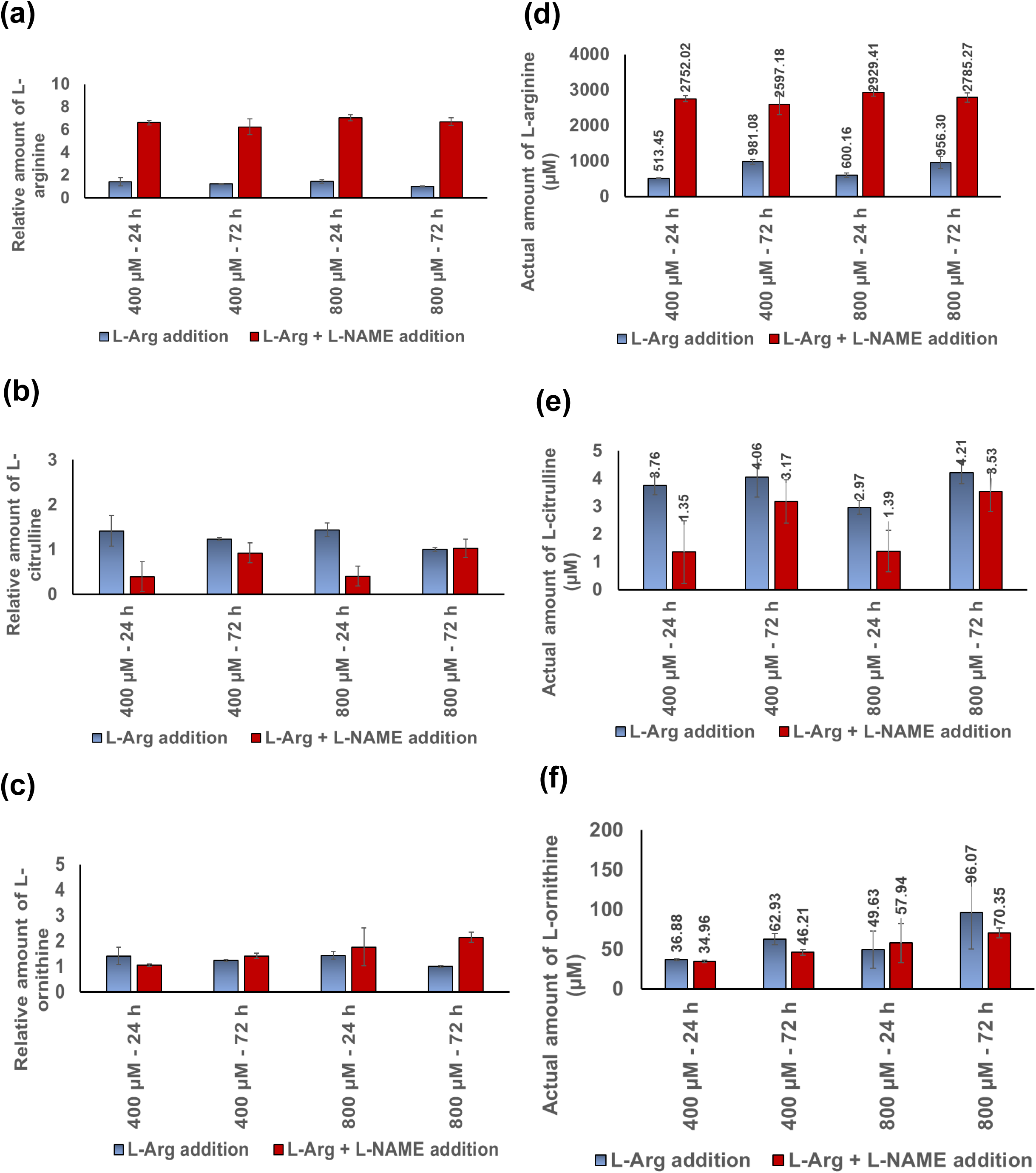
Residual serum L-Arg, L-Cit and L-Orn in BNL CL2 cells with exogenous L-Arg and L-NAME (4 mM). Relative and actual residue amount of L-Arg **(a and b)**, L-Cit **(c and d)** and L-Orn **(e and f)** analysed in plasma samples of BNL CL2 cells cultured with or without the presence of iNOS inhibitor; L-NAME (4 mM) in excess amount of L-Arg (400 and 800 µM) after 24 and 72 h. Data points represent the mean ± SD of each sample. Error bars represent the standard deviation from the mean (n =3).

The amino acid synthesised via L-Arg/Arginase pathway, L-ornithine (Figures 9c **and 9f**) had decreased (P > 0.05) in the samples cultured with 400 µM L-Arg and L-NAME (0.96-fold) at 24 h, whereas it had increased (P > 0.05) with 800 µM L-Arg with L-NAME (1.36-fold) compared to the samples cultured without L-NAME at the same time point. Interestingly, the amount of culture supernatant L-ornithine was increased (P<0.001) with culture time when the cells in 400 (1.32-fold) and 800 µM L-Arg (1.21-fold) were treated with L-NAME.

The amount of L-Arg present in the samples with SNAP was investigated and is presented in (**Figures 10a and 10d**). The level of cell culture supernatant L-Arg in the samples cultured in excess exogenous L-Arg (400 and 800 µM) and treated with SNAP was decreased (P<0.0001) after 6 (400 µM + SNAP; 0.84-fold and 800 µM + SNAP; 0.72-fold) and 24 h (400 µM + SNAP; 0.41-fold and 800 µM + SNAP; 0.82-fold) compared to the samples without SNAP at the same time points.

**Figure 10.**
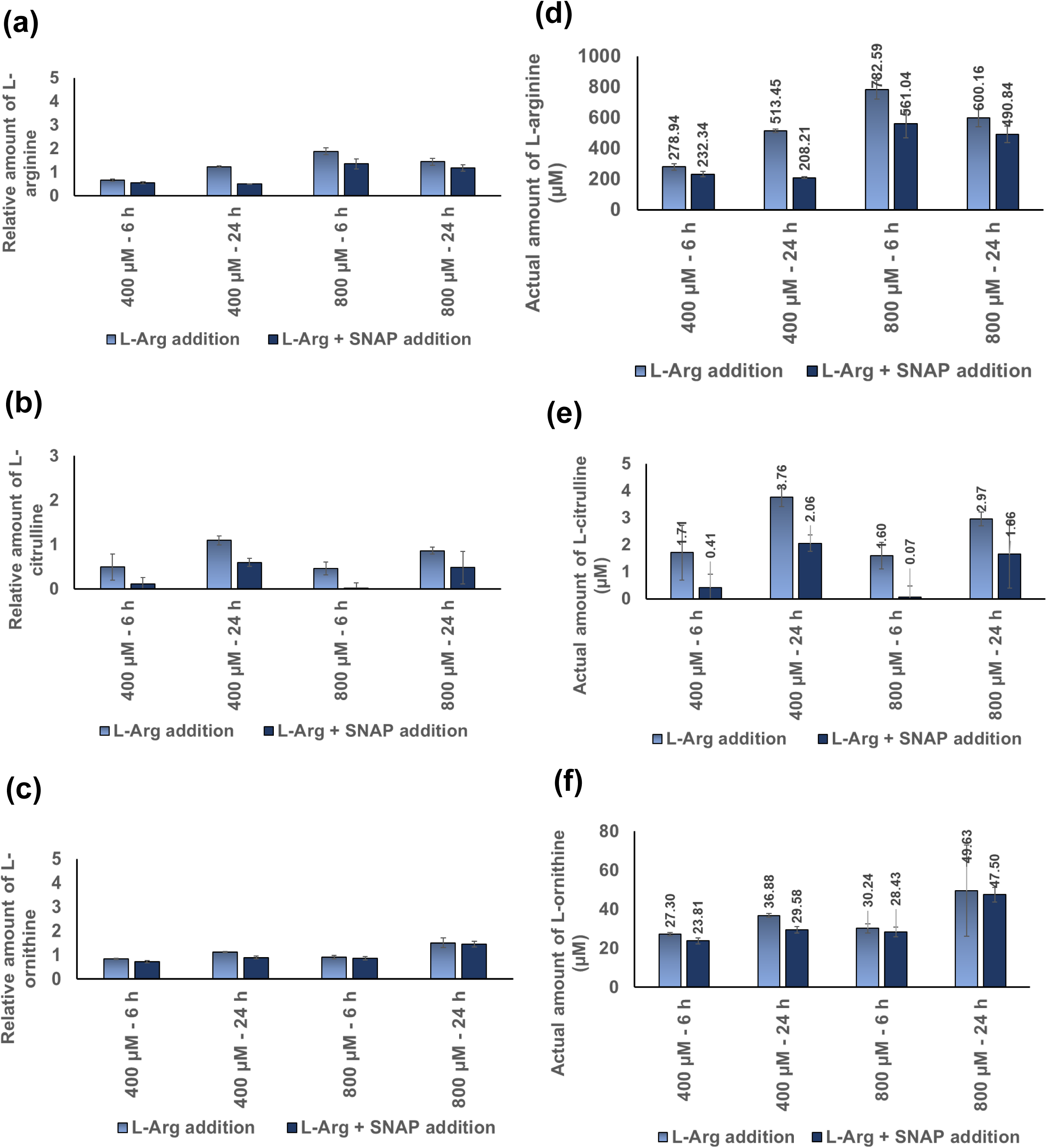
Residual serum L-Arg, L-Cit and L-Orn in BNL CL2 cells with exogenous L-Arg and SNAP (100 µM). Relative and actual residue amount of L-Arg **(a and b)**, L-Cit **(c and d)** and L-Orn **(e and f)** analysed in plasma samples of BNL CL2 cells cultured with or without the presence of external NO donor; SNAP (100 µM) in excess amount of L-Arg (400 and 800 µM) after 6 and 24 h. Data points represent the mean ± SD of each sample. Error bars represent the standard deviation from the mean (n =3).

L-citrulline concentrations upon SNAP addition is also shown in the (**Figures 10b and 10e**). SNAP addition to cells cultured in L-arginine (400 and 800 µM) reduced (P<0.0001) the level of L-citrulline present after 6 (400 µM + SNAP; 0.24-fold and 800 µM + SNAP; 0.043-fold) and 24 h (400 µM + SNAP; 0.55-fold and 800 µM + SNAP; 0.56-fold) compared to the samples cultured without SNAP at the same time points.

Finally, the amount of L-ornithine in the cell culture supernatant samples cultured with L-Arg and SNAP was analysed and is presented in **Figures 10c and 10f**). As with the other amino acids monitored, L-Arg and L-Cit, overall, the amount of L-ornithine was decreased in the samples cultured with SNAP but not to a significant amount (P=0.3062).

## Discussion

In this study, the impact of excess exogenous L-Arg in the media of and *in-vitro* cultured model cell line and the subsequent impact on NO signalling and L-Arg metabolism has been investigated. The cellular response of BNL CL2 cells growth in different exogenous L-arginine concentrations has been evaluated. BNL CL2 cells are derived from a mouse liver and are thus an *in-vitro* cell culture model to investigate how liver cells respond to differing exogenous L-arginine concentrations. The liver plays an important role in the metabolism of L-arginine [20]. Previous studies have reported that the ‘average’ plasma concentration of L-arginine in fed-rats is 175 µM [5] and therefore the initial concentrations investigated here were at a similar and higher range. Other studies have indicated that there are no adverse effects in experimental adult rats chronically administered, via enteral diets, large amounts of L-arginine (2.14 g/kg body weight-1 d-1) [21,22]. However, a recent review article suggests that L-arginine supplementation to different cell models within the range of 100 µM – 100 mM reflects the impact of L-arginine in the treatment of carbohydrate and lipid metabolism disorders [23]. Therefore, the concentrations of arginine used in this study (400 to 800 µM) are relevant to the physiological ranges found in mammals and to the nutritional and clinical concentrations impacting health and disease in mammals. It is important to note that in most cases the most prevalent impacts were observed with 800 µM culturing.

Studies have shown that increasing extracellular concentrations of L-arginine results in an increase in the intracellular concentration of L-arginine in a dose dependent manner. For example, an *in-vitro* study in bovine aortic endothelial cells cultured in varying extracellular concentrations of arginine (0.1 to 10 mM) confirmed that increasing extracellular L-arginine produced a dose-dependent increase in intracellular arginine [24]. In line with these findings, we hypothesised that exogenous L-arginine supplementation (400 and 800 µM) of BNL CL2 cultured cells would impact intracellular signalling of L-arginine responsive pathways. To determine if this was correct, gene and protein expression of L-arginine responsive pathways were examined to explore the molecular mechanisms and cellular responses in BNL CL2 cells as a model.

In order to determine the impact of different exogenous L-arginine concentrations on the liver derived BNL CL2 cell line, target mRNA, protein and protein post-translational modifications were monitored in L-arginine responsive pathways. The key downstream target of the L-arginine/NO metabolic pathway, AMPK, increased at the transcript level (RE 3.29; P < 0.0001) when cells were cultured in L-arginine concentrations of 800 µM compared to that observed in control complete DMEM media cultured samples at 24 h. The expression of AMPK is known to have specificity to tissue and localization within cells [25]. Interestingly, the profile of the impact of additional exogenous L-arginine concentrations on AMPK expression at the transcript (mRNA) and protein level was generally similar but not the same with large difference at 400 µM after 72 h and 800 µM at 24 h (Figures 1c **and 4a**). This may reflect a time dependent link between changes in transcription before changes in protein synthesis are observed and the fact that there are a number of metabolic signalling pathways that converge and cross-talk with AMPK to coordinate responses to nutritional and hormonal signalling of the cells [26]. With regard to time dependent changes, it would be interesting in future studies to determine if the drop in transcript amounts in 400 µM cultures at 72 h resulted in a drop in protein amounts at a later time point for example. Further, in this study AMPK transcript expression was analysed by monitoring one of the catalytic subunits of AMPK isoforms; AMPKα1.

However, in protein analysis, the primary antibody for anti-AMPK detects both isoforms of AMPKα (AMPKα1 and AMPKα2) but does not distinguish the isoforms present in the samples. This may also account for differences between the protein and transcript data.

In addition to AMPK protein amounts, phosphorylation was also investigated. Phosphorylated AMPK levels were increased in response to increased extracellular concentrations of L-arginine from 0 to 400 and 800 μM in a culture time dependent manner. Phosphorylation of AMPK activates AMPK and the phosphorylation status thus controls overall cellular lipid metabolism [27]. Activated AMPK phosphorylates its downstream target ACC-1 [28] and in this study the levels of phosphorylated ACC-1 in 800 µM L-arginine cultured samples at 72 h were increased by 6-fold over that of the control complete media DMEM. Phosphorylation of AMPK inactivates ACC-1 enzyme and suppresses fatty acid synthesis. This is a key mechanism by which AMPK responds to low energy states by turning off energy-consuming processes and promoting energy-producing ones. If this was the case it would be expected that the inactivation of ACC-1 via its phosphorylation would lead to an increase in the activity of CPT-1, facilitating the transport of long-chain fatty acids from the cytosol to the mitochondria for oxidation [29,30]. However, in the BNL CL2 cells, the concentrations of L-arginine investigated did not affect CPT-1 gene expression at the mRNA transcript level and consequently there was no difference in CPT-1A protein expression in 800 µM L-arginine cultures at 72 h compared with the control. However, perhaps unexpectedly, in L-arginine cultures at 400 µM at 72 h there was an increased level of CPT-1A protein (2-fold) in comparison to the control complete DMEM cultures.

When the lipogenic genes ACC-1 (RE 7.46) and SREBP-1 (RE 6.76) were investigated the expression increased at the mRNA level, whereas the AMPK mRNA was lowest, in cells cultured in 400 mM L-Arg for 72 h in comparison with the complete DMEM control. The highest mRNA amounts of these lipogenic mRNAs were accompanied by increases in the respective proteins at the same time point;

ACC-1 protein expression was 16-fold higher than the control and the catalytically active form of SREBP-1 was increased (1.7-fold) at the same time point. This suggests an inverse relationship between AMPK and expression of these proteins responsible for the induction of lipogenesis by the liver. When BNL CL2 cells treated with L-NAME, AMPK transcript was decreased but there was increased ACC-1 mRNA expression. The investigation of the effect and mechanism of action of the NOS inhibitor L-NAME on the expression of AMPK and ACC-1 in BNL CL2 cell revealed that an inhibition of NO synthesis moderately attenuated the effect of L-arginine on the expression of AMPK (and ACC-1) at the mRNA and protein level, but greatly impacted the amount of nitrite in the cell culture supernatant (Figure 7) and residual amino acids in BNL CL2 (Figure 9). The proposed mechanisms and cellular responses by which liver cells (BNL CL2) respond to excess exogenous L-Arg (400 and 800 µM) and its modulators (L-NAME and SNAP) are summarised in Figure 11. In summary, additional L-arginine supplementation increased expression of key enzymes for fatty acid oxidation in the liver, while decreasing hepatic expression of key enzymes for lipogenesis and cholesterol synthesis. These coordinated changes in gene and protein expression among insulin-sensitive tissue may partially provide a molecular mechanism for the fat mass deposition during obesity and how that would be overcome by L-Arg supplementation.

**Figure 11.**
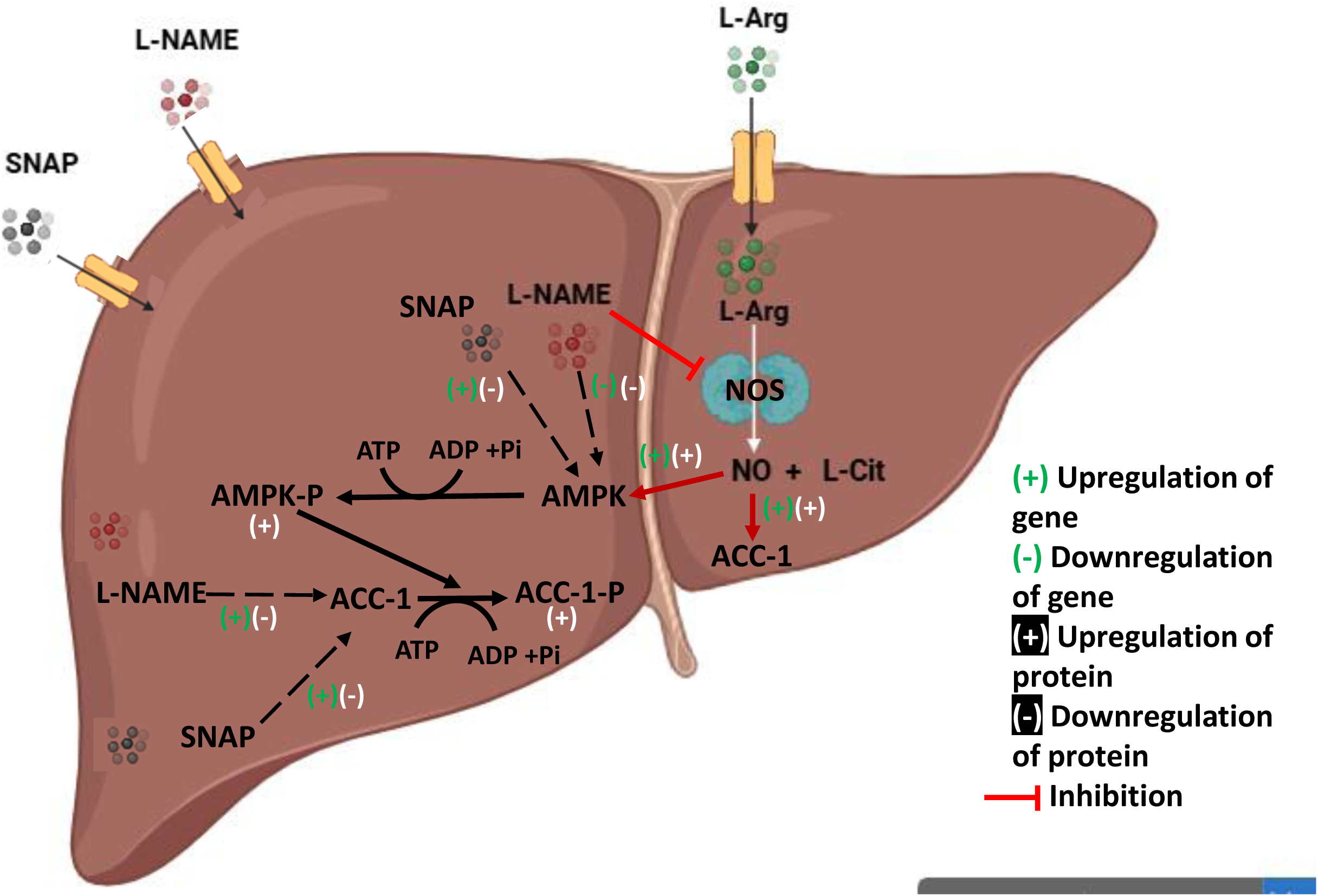
Putative cell signalling pathway of L-Arg/NOS/NO and regulation of this pathway by the modulators. The proposed mechanisms responsible for the beneficial effect of L-Arg/NOS/NO on the metabolic pathways in mammalian liver cells, BNL CL2. The symbol (+) denotes an upregulation in gene expression or protein expression. The symbol (-) denotes a downregulation of gene expression or protein expression.

## Materials and Methods

### Establishment of cell culture models for investigating cell fitness and AMPK and ACC-1 cell signalling upon addition of exogenous L-Arg

#### Preparation of stock solutions of L-Arg and growth medium with varying L-Arg concentrations

Growth medium with different L-Arg concentrations was created using commercially available DMEM media for Stable Isotope Labelling using Amino Acids (SILAC DMEM, Thermo Scientific, United States), which is lacking in both L-lysine and L-Arg. Foetal bovine serum (FBS, Sigma, United States), was added to a final concentration of 10% (v/v). L-lysine-HCl (Sigma, United States) was also added to give the same concentration of L-lysine (146 mg/L) as in the complete DMEM basal media formula described in Thermo Fisher Scientific company (DMEM, Cat. No: 41966, United States). This media was referred to as L-Arg deficient media since it did not contain any exogenously added L-Arg. Although FBS may contain unknown concentrations of L-Arg, the quantity of L-Arg in the media was found to be undetectable by HPLC following the addition of FBS, hence this media was deemed L-Arg deficient (0 µM). To create an L-Arg stock solution (71.75 mM), L-Arg (Sigma, United States, 0.125 g) was added to 10 mL of SILAC L-Arg deficient DMEM medium. To create various L-Arg concentration-containing media, this was subsequently diluted as needed using the SILAC media containing L-Lys and FBS.

#### Cell culture

The BNL CL2 cell line was used as a model system, which was a generous gift from Associate Prof of Medicine Dimiter Avtanski, Zucker School of Medicine at Hofstra/Northwell, Hempstead, New York. The BNL CL2 cell line is a mouse hepatocyte epithelial insulin-sensitive cell [31–33]. BNL CL2 cells were seeded into 6-well tissue culture plates at 2×10^5^ viable cells/well in 2 mL of complete DMEM media and then incubated in a static incubator for 24 h at 37°C, 5% CO_2_. After 24 h, the media was replaced with media of different concentrations of L-Arg (0, 400 and 800 µM) or the control complete DMEM with 10% (v/v) FBS (2 mL/ well). Complete DMEM is meant to contain 398 µM L-Arg of L-Arg hydrochloride according to the media formulation from the company (Thermo Fisher, United States), however the concentration was determined as 250 µM by the HPLC method used in this study. In comparison to the full DMEM controls, the exogenous L-Arg concentration increased 1.6 and 3.2 times under the 400 and 800 µM culture conditions. Thus, the 400 and 800 µM culture conditions represent a 1.6 and 3.2-fold increase in exogenous L-Arg concentration compared to the complete DMEM controls. Cells were then harvested at time points; 24, 48, 72 and 120 h to measure culture viability (%) and the viable number of cells/mL (x10^6^) using a Vi-CELL cell viability analyzer (Beckman Coulter, Life Sciences, United States). In order to examine the expression of mRNA transcripts and proteins, and measure L-Arg metabolites in the cell culture supernatant, the cells and supernatant were also collected 24 and 72 hours after the addition of L-Arg medium at varying concentrations. The BNL CL2 cells and cell culture supernatant were also harvested at T=0; untreated samples (24 h after incubation of the cells and collected before L-Arg addition).

In some experiments, after 24 h, the media was replaced with media with different concentrations of L-Arg (0, 400 and 800 µM) or the control complete DMEM with 10% (v/v) FBS (2 mL/ well) with additions of L-NAME (a NOS inhibitor, 4 mM) and cells and cell culture supernatant then harvested at two time points; 24 and 72 h post addition. In another set of experiments, after 24 h the media was replaced with media of different concentrations of L-Arg (0, 400 and 800 µM) or the control complete DMEM with 10% (v/v) FBS (2 mL/ well) with addition of SNAP (a NO donor, 100 µM) and cells and cell culture supernatant harvested 6 and 24 h post addition.

Subsequently samples were lysed for mRNA or protein analysis. The cell culture media (2 mL) from additional experiments was also collected from the cell cultures and frozen at-20°C.

#### Quantitative real-time PCR (qRT-PCR) for transcript mRNA analysis

BNL CL2 cells were lysed using RLT (RNA lysis buffer, 350 µL, Qiagen, Germany). Collected RNA lysates were homogenised using a QIAshredder kit (Qiagen, Germany). The commercially available RNeasy Mini Kit (Qiagen, Germany) was then used to extract total RNA in accordance with the manufacturer’s instructions. The RQ1 RNase-Free DNase kit (Promega, USA) was used to treat extracted total RNA for contaminating DNAase.

Primers (described in **Table S1** in Supplementary Materials) were designed to target genes of interest. qRT-PCR amplification of target sequences and the housekeeping control (β-actin) was undertaken using the commercially available iTaqTM Universal SYBR Green One-Step Kit (Bio-Rad, United States). The amplicons were amplified and their melting curve profiles were created using a DNA engine opticon 2 system for real-time PCR detection thermocycler (Bio-Rad, USA).

#### Protein and post-translational modification (phosphorylation) analysis using western blotting

Samples for protein analysis were lysed with lysis buffer (200 µL/well). Protein content in samples was determined using the Bradford assay. Samples were prepared with x 5 sample/loading buffer (reducing buffer). The ratio between reducing sample buffer (x 5): diluted sample was 1:5. The required volume of protein extract was diluted with water to obtain x 1 final sample buffer concentration with 10 and 20 µg of protein/25 µL total sample volume/lane. The proteins were separated using 10% SDS-PAGE. Wet transfer conditions were then used to transfer the protein onto nitrocellulose blotting membranes. The proteins were incubated with appropriate primary antibody (detailed in **Table S2** in Supplementary Materials) overnight at 4°C. Then, using an Optimax 2010 film processor (PROTEC GmbH & Co, Germany), the blot was probed with a suitable secondary antibody linked to horseradish peroxidase for one hour at room temperature for development.

Densitometry of bands on western blots was undertaken using the open software package ImageJ (National Institutes for Health (NIH), USA).

#### Nitric oxide / Nitrite measurement by Griess Assay

After collecting 2 mL of frozen cell culture media from L-Arg and/or L-NAME and SNAP additions to BNL CL2 cells, the supernatant without cell pellet was collected after vortexing and centrifuging for 10 minutes at 1500 rpm. Cell culture supernatant (50 µL) was used in the assay. Using sodium nitrite as a reference, the Griess test [33,34] was used to measure the amount of nitrite accumulated in cell culture supernatants following L-Arg supplementation.

#### Analysis of L-arginine, L-ornithine and L-citrulline by HPLC

The cell free culture supernatant for HPLC analysis was obtained by vortexing and centrifuging the collected serum samples for 10 minutes at 1500 rpm. Samples were deproteinised using a modified protocol of that described by [35]. After adding 100% (v/v) ice-cold ethanol (400 µL) to the cell culture supernatant (100 µL), the mixture was aggressively vortexed for 15 minutes at room temperature in order to extract FBS and cellular proteins. Amino acids were derivatized at room temperature using a pre-column derivatisation method. A pre-column derivatization agent, O-phthaldialdehyde (OPA) reagent complete solution consisting OPA (Sigma, USA, 1 mg/mL), specifically designed for primary amines and amino acids at alkaline pH was used. Mobile phase A (0.1 M sodium acetate, pH 7.2) was prepared as described by (Wu and Meininger 2008). The mobile phase B was 100% (v/v) methanol. A standard amino acid mixture (6 mM), consisting of the essential 20 amino acids, was a gift from Dr Andrew Lawrence, School of Biosciences, University of Kent, and was prepared as outlined previously [37]. Furthermore, to form an extended amino acid standard solution, L-citrulline (Sigma, USA) and L-ornithine monohydrochloride (Sigma, USA) (0.6 mM of each) were added to the standard amino acid mixture from the respective stock solutions of L-citrulline and L-ornithine (6 mM of each). To a dark glass vial (1.5 mL), an extended amino acid standard mixture (0.6 mM each) or test cell culture sample solution (100 µL) was added. An Agilent 1100 HPLC (Agilent Technologies, Germany) equipped with a diode-array detector (DAD) was used. Samples were injected (25 μL) onto an ACE HPLC column; RP-C18, dimensions 125 × 4.6 mm, 5 μm (Avantor, United States). 25 µL of a standard or sample solution and 25 µL of the OPA reagent solution were combined by the autosampler automatically, and the mixture was then incubated for one minute in a reaction loop. The autosampler was automated to mix 25 µL of a standard or sample solution with 25 µL of the OPA reagent solution and allowed for incubation for 1 min in a reaction loop. Amino acids were separated with a linear gradient (**Table S3** in Supplementary Materials) with a total running time of 49 min; flow rate, 1.1 mL/min. The detector, Diode Array Detector (DAD), was programmed to switch to 338 nm, 10 nm bandwidth, and reference wavelength 390 nm, 20 nm bandwidth.

The molar absorptivity of each derivatized amino acid was detected at 338 nm (λmax).

The stock solutions of L-Arg, L-Cit, and L-Orn were used to generate the standard curves for L-Arg, L-Cit and L-Orn. Identification of particular amino acid signals was based on the comparison between the retention time of the extended amino acid standard mixture and the amino acids of interest from analysis of samples.

Quantitation was based on the standard curve method using a linear curve fitted by linear regression analysis. The equation of the line fitted for the standard curves of amino acids was used to determine the unknown concentration of each amino acid in experimental samples.

## Statistical analysis

Statistical analysis was undertaken using GraphPad Prism 9.4.1 software for Windows (GraphPad Software, San Diego California USA) and Microsoft Excel. Samples were analysed in triplicate biological replicates. Two-way ANOVA was used to analyze the means and standard deviation of the data. The means of the treatment groups (0, 400, and 800 µM L-Arg) were compared to the control complete DMEM media addition at time points 24 and 72 hours using the Tukey’s multiple comparison method. To compare between multiple treatments groups, the Bonferroni’s multiple comparison method was used to determine differences among the means of the treatment groups (0, 400 and 800 µM L-Arg) and the control complete DMEM media addition with the addition of either L-NAME or SNAP in BNL CL2 cells across the time points either 24 and 72 h or 6 and 24 h. Statistical significance was considered as probability values ≤ 0.05.

## Ethics approval and consent to participate

Not applicable.

## Consent for publication

Not applicable.

## Financial interests

The author has no relevant financial or non-financial interests to disclose.

## Authors’ contributions

SP participated in the conception and design of the experiments under the guidance of CMS; SP performed all of the experiments under the supervision of CMS; SP analyzed and interpreted the data; SP and CMS wrote the paper. The authors read and approved the final manuscript.

## Competing interests

The authors declare that they have no conflict of interest.

## Funding

This research was partially supported by a grant ‘Accelerating Higher Education and Development Program (AHEAD) Operation, a World Bank funded project to accelerate higher education expansion and development in Sri Lanka (AHEAD/PhD/R1/AH/036).

## Supporting information

Statistical Analysis Tables

## Acknowledgements

We thank Dr. Mark Shepherd for assistance with the Griess assay. Thanks to Dr. Andrew Lawrence, who helped with the HPLC analysis, as well as Dr. Dave Beal, who helped with the LC-MS experiment for analyses of metabolites.

## Data availability statement

The data supporting the findings of this study have been included as part of the Supplementary Materials. Also, the statistical tables are provided in the Supplementary Materials. No additional datasets were generated or analysed during this study.

## Electronic supplementary materials

Supplementary materials

Statistical analysis tables

## Abbreviations

BNL CL2: Mouse hepatocytes spontaneously immortalized cell line
ACC: Acetyl-CoA carboxylase
AMPK: 5’-Adenosine monophosphate-activated protein kinase
HPLC: High-performance liquid chromatography
L-Arg: L-arginine
L-Cit: L-citrulline
L-NAME: N^G^-nitro-L-Arg methylester
L-Orn: L-ornithine
NOS: Nitric oxide synthase
NO: Nitric oxide
NO_2_^-^: Nitrite
SILAC DMEM: Stable isotope labeling with amino acids in cell culture DMEM
SNAP: S-nitroso-N-acetyl-DL-penicillamine

## Notes

### Competing Interest Statement

The authors have declared no competing interest.

