## Supplementary material for "L-Arginine supplementation modulates L-Arg/NO metabolic processes and AMPK/ACC-1 signalling in BNL CL2 hepatocytes": Statistical Analysis Tables

**Table 1** Statistical table for cell growth profile of BNL CL2 cells with different concentrations of L-arginine and the control.

| Source of Variation | P value | P value summary | Significant? |
|---|---|---|---|
| **Interaction** | <0.0001 | **** | Yes |
| **Time point** | <0.0001 | **** | Yes |
| **L-Arg +/- and Control (Com)** | <0.0001 | **** | Yes |

**Table 2** Statistical table for culture viability profile of BNL CL2 cells grown in different concentrations of L-arginine and the control.

| Source of Variation | P value | P value summary | Significant? |
|---|---|---|---|
| **Interaction** | 0.0225 | * | Yes |
| **Time point** | 0.7313 | ns | No |
| **L-Arg +/- and Control (Com)** | 0.0766 | ns | No |

**Table 3** Samples comparison for cell growth profile and culture viability profile of BNL CL2 cells grown in different concentrations of L-arginine and the control.

| Samples comparison Tukey multiple comparison | Variables: Cell fitness parameters | | | | | | | |
|---|---|---|---|---|---|---|---|---|
| | Cell growth | | | | Culture viability | | | |
| | T=24 h | T=48 h | T=72 h | T=120 h | T=24 h | T=48 h | T=72 h | T=120 h |
| **Cont. Com vs. 400 µM** | ns | * | ns | *** | ns | ns | ns | ns |
| **Cont. Com vs. 800 µM** | ns | ** | ns | **** | ns | ** | ns | ns |
| **Cont. Com vs. No L-Arg** | ns | **** | **** | **** | ns | ns | ns | ns |

**Tables 1-3** Statistical tables for cell growth/viable cell number and culture viability profiles of BNL CL2 cells with different concentrations of L-arginine (0, 400 and 800 µM) and the control complete DMEM media at 24, 48, 72 and 120 h. The tables summarise two-way ANOVA followed by a Tukey multiple comparison test using GraphPad Prism 9.4.1. The stars flag the levels of significance; ns = $P > 0.05$, * = $P \leq 0.05$, ** = $P \leq 0.01$, *** = $P \leq 0.001$ and **** = $P \leq 0.0001$.

**Table 4** Statistical table for relative mRNA transcript expression (ΔΔCt) of AMPK and ACC-1 upon L-Arg addition

| | AMPK | | ACC-1 | |
|---|---|---|---|---|
| **Source of Variation** | **P value** | **Significant?** | **P value** | **Significant?** |
| **Interaction** | **** | Yes | **** | Yes |
| **Time point** | **** | Yes | **** | Yes |
| **L-Arg +/- and control (Com)** | **** | Yes | **** | Yes |

**Table 5** Samples comparison of relative mRNA transcript expression (ΔΔCt) of AMPK and ACC-1 upon L-Arg addition

| Samples comparison Tukey multiple comparison | Variables: Gene expression | | | | | |
|---|---|---|---|---|---|---|
| | AMPK | | | ACC-1 | | |
| | T=0 | T=24 h | T=72 h | T=0 | T=24 h | T=72 h |
| **Cont. Com vs. untreated** | ** | *** | **** | ns | ns | ns |

| | | | | | | |
|---|---|---|---|---|---|---|
| **Cont. Com vs. 400 µM** | ns | ns | *** | ns | ns | **** |
| **Cont. Com vs. 800 µM** | ns | **** | ** | ns | ns | ns |
| **Cont. Com vs. No L-Arg** | ns | ns | ** | ns | ns | **** |

**Tables 4-5** Statistical tables for relative mRNA transcript expression (ΔΔCt) of AMPK and ACC-1 in BNL CL2 cells cultured in medium of different L-arginine concentrations (0, 400 and 800 µM) and control complete DMEM media 24 and 72 h after addition. Untreated cultures at T=0. Tables summarise two-way ANOVA followed by a Tukey multiple comparison test using GraphPad Prism 9.4.1. The stars indicate the levels of significance; ns = $P > 0.05$, * = $P \leq 0.05$, ** = $P \leq 0.01$, *** = $P \leq 0.001$ and **** = $P \leq 0.0001$.

**Table 6** Statistical table for relative mRNA transcript expression (ΔΔCt) of AMPK and ACC-1 upon L-Arg addition with L-NAME

| | **AMPK** | | **ACC-1** | |
|---|---|---|---|---|
| **Source of Variation** | **P value** | **Significant?** | **P value** | **Significant?** |
| **Interaction** | **** | Yes | ns | No |
| **Samples, conditions and time points** | *** | Yes | **** | Yes |
| **L-Arg +/- L-NAME** | **** | Yes | ns | No |

**Table 7** Samples comparison of relative mRNA transcript expression (ΔΔCt) of AMPK and ACC-1 upon L-Arg addition with L-NAME

| **Samples comparison Bonferroni's multiple comparisons** | **L-Arg addition and L-Arg + L-NAME addition** | |
|---|---|---|
| | **AMPK** | **ACC-1** |
| **Cont. Com - 24 h** | ns | ns |
| **Cont. Com - 72 h** | ns | ns |
| **400 µM - 24 h** | ns | ns |
| **400 µM - 72 h** | * | ns |
| **800 µM - 24 h** | ns | ns |
| **800 µM - 72 h** | ns | ns |
| **No L-Arg - 24 h** | * | ns |
| **No L-Arg - 72 h** | **** | ns |

**Table 8** Statistical table for relative mRNA transcript expression (ΔΔCt) of AMPK and ACC-1 upon L-Arg addition with SNAP

| | AMPK | | ACC-1 | |
|---|---|---|---|---|
| **Source of Variation** | **P value** | **Significant?** | **P value** | **Significant?** |
| **Interaction** | ** | Yes | *** | Yes |
| **Samples, conditions and time points** | ns | No | ns | No |
| **L-Arg +/- SNAP** | **** | Yes | **** | Yes |

**Table 9** Samples comparison of relative mRNA transcript expression (ΔΔCt) of AMPK and ACC-1 upon L-Arg addition with SNAP

| **Samples comparison Bonferroni's multiple comparisons** | **L-Arg addition and L-Arg + SNAP addition** | |
|---|---|---|
| | **AMPK** | **ACC-1** |
| **Cont. Com - 6 h** | *** | * |
| **Cont. Com - 24 h** | ns | *** |
| **400 µM - 6 h** | * | ns |
| **400 µM - 24 h** | ns | ns |
| **800 µM - 6 h** | ns | ns |
| **800 µM - 24 h** | ns | ns |
| **No L-Arg - 6 h** | ** | *** |
| **No L-Arg - 24 h** | * | **** |

**Tables 6-9** Statistical tables for relative mRNA transcript expression (ΔΔCt) of AMPK and ACC-1 in BNL CL2 cells cultured in medium of different L-arginine concentrations (0, 400 and 800 µM) and control complete DMEM media with nitric oxide synthase inhibitor; L-NAME (4 mM) for 24 and 72 h after addition and the NO donor SNAP (100 µM) for 6 and 24 h. The tables summarise two-way ANOVA followed by a Bonferroni multiple comparison test using GraphPad Prism 9.4.1. The stars indicate the levels of significance; ns = P > 0.05, * = P ≤ 0.05, ** = P ≤ 0.01, *** = P ≤ 0.001 and **** = P ≤ 0.0001.

**Table 10** Statistical table for relative protein amounts for total AMPKα and phosphorylated AMPKα at Thr172 (AMPKα-P) upon L-Arg addition

| | AMPK | | AMPK-P | |
|---|---|---|---|---|
| **Source of Variation** | **P value** | **Significant?** | **P value** | **Significant?** |
| **Interaction** | **** | Yes | **** | Yes |
| **Time** | **** | Yes | **** | Yes |
| **L-Arg +/- and control (Com)** | **** | Yes | **** | Yes |

**Table 11** Statistical table for relative protein amounts for total ACC-1 and phosphorylated ACC-1 at Ser79 (ACC-1-P) upon L-Arg addition

| | ACC-1 | | ACC-1-P | |
|---|---|---|---|---|
| **Source of Variation** | **P value** | **Significant?** | **P value** | **Significant?** |
| **Interaction** | **** | Yes | **** | Yes |
| **Time** | **** | Yes | **** | Yes |
| **L-Arg +/- and control (Com)** | **** | Yes | **** | Yes |

**Table 12** Samples comparison of relative protein amounts for total AMPKα and ACC-1 and phosphorylated AMPKα at Thr172 (AMPKα-P) and ACC-1 at Ser79 (ACC-1-P) upon L-Arg addition

| Samples comparison Tukey multiple comparison | Variables: Protein expression | | | | | | | | | | | |
|---|---|---|---|---|---|---|---|---|---|---|---|---|
| | AMPK | | | AMPK-P | | | ACC-1 | | | ACC-1-P | | |
| | T=0 | T=24 h | T=72 h | T=0 | T=24 h | T=72 h | T=0 | T=24 h | T=72 h | T=0 | T=24 h | T=72 h |
| **Cont. Com vs. untreated** | ns | ns | **** | **** | ns | **** | ns | ns | ns | *** | ns | ** |
| **Cont. Com vs. 400 µM** | ns | ns | * | ns | ns | **** | ns | **** | **** | ns | **** | **** |
| **Cont. Com vs. 800 µM** | ns | ns | ns | ns | ns | **** | ns | **** | **** | ns | **** | **** |
| **Cont. Com vs. No L-Arg** | ns | ns | ** | ns | **** | ns | ns | ns | ns | ns | ns | ns |

**Tables 10-12** Statistical tables for relative protein amounts for total AMPKα and ACC-1 and phosphorylated AMPKα at Thr172 (AMPKα-P) and ACC-1 at Ser79 (ACC-1-P) in BNL CL2 cells. The tables summarise two-way ANOVA followed by a Tukey multiple comparison test using GraphPad Prism 9.4.1. The stars indicate the levels of significance; ns = P > 0.05, * = P ≤ 0.05, ** = P ≤ 0.01, *** = P ≤ 0.001 and **** = P ≤ 0.0001.

**Table 13** Statistical table for relative protein amounts for total AMPKα and ACC-1 upon L-Arg with L-NAME addition

| Source of Variation | AMPK | | | ACC-1 | | |
|---|---|---|---|---|---|---|
| | P value | P value summary | Significant? | P value | P value summary | Significant? |
| Interaction | <0.0001 | **** | Yes | <0.0001 | **** | Yes |
| Samples, conditions and time points | 0.0044 | ** | Yes | <0.0001 | **** | Yes |
| L-Arg +/- L-NAME | <0.0001 | **** | Yes | <0.0001 | **** | Yes |

**Table 14** Samples comparison of relative protein amounts for total AMPKα and ACC-1 upon L-Arg with L-NAME addition

| Samples comparison Bonferroni's multiple comparisons | Variables: Protein expression L-Arg addition and L-Arg + L-NAME addition | |
|---|---|---|
| | AMPK | ACC-1 |
| Cont. Com - 24 h | ns | **** |
| Cont. Com - 72 h | ** | **** |
| 400 µM - 24 h | * | **** |
| 400 µM - 72 h | **** | **** |
| 800 µM - 24 h | * | **** |
| 800 µM - 72 h | ** | **** |
| No L-Arg - 24 h | ns | *** |
| No L-Arg - 72 h | ns | ** |

**Table 15** Statistical table for relative protein amounts for total AMPKα and ACC-1 upon L-Arg with SNAP addition

| Source of Variation | AMPK | | | ACC-1 | | |
|---|---|---|---|---|---|---|
| | P value | P value summary | Significant? | P value | P value summary | Significant? |
| **Interaction** | 0.2117 | ns | No | <0.0001 | **** | Yes |
| **Samples, conditions and time points** | 0.0584 | ns | No | <0.0001 | **** | Yes |
| **L-Arg +/- SNAP** | <0.0001 | **** | Yes | <0.0001 | **** | Yes |

**Table 16** Samples comparison of relative protein amounts for total AMPKα and ACC-1 upon L-Arg with SNAP addition

| Samples comparison Bonferroni's multiple comparisons | Variables: Protein expression L-Arg addition and L-Arg + SNAP addition | |
|---|---|---|
| | **AMPK** | **ACC-1** |
| **Cont. Com - 24 h** | ns | ns |
| **400 µM - 24 h** | ** | **** |
| **800 µM - 24 h** | ** | **** |
| **No L-Arg - 24 h** | * | ns |

**Tables 13-16** Statistical tables for relative mRNA transcript expression (ΔΔCt) of AMPK and ACC-1 in BNL CL2 cells cultured in medium of different L-arginine concentrations (0, 400 and 800 µM) and control complete DMEM media with addition of either L-NAME (4mM) or SNAP (100 µM) in BNL CL2 cells across the time points either 24 and 72 h or 6 and 24 h. The tables summarise two-way ANOVA followed by a Bonferroni's multiple comparisons test using GraphPad Prism 9.4.1. The stars indicate the levels of significance; ns = P > 0.05, * = P ≤ 0.05, ** = P ≤ 0.01, *** = P ≤ 0.001 and **** = P ≤ 0.0001.

**Table 17** Statistical table for the effect of exogenous L-arginine concentration on nitrite production**.**

| Source of Variation | P value | P value summary | Significant? |
|---|---|---|---|
| **Interaction** | <0.0001 | **** | Yes |
| **Time** | <0.0001 | **** | Yes |
| **L-Arg +/- and Control (Com)** | 0.0002 | *** | Yes |

**Table 18** Statistical table for the effect of exogenous L-arginine concentration on quantification of residual serum L-Arg.

| Source of Variation | P value | P value summary | Significant? |
|---|---|---|---|
| **Interaction** | <0.0001 | **** | Yes |
| **Time** | <0.0001 | **** | Yes |
| **L-Arg +/- and Control (Com)** | <0.0001 | **** | Yes |

**Table 19** Statistical table for the effect of exogenous L-arginine concentration on quantification of residual serum L-Cit.

| Source of Variation | P value | P value summary | Significant? |
|---|---|---|---|
| **Interaction** | <0.0001 | **** | Yes |
| **Time** | <0.0001 | **** | Yes |
| **L-Arg +/- and Control (Com)** | <0.0001 | **** | Yes |

**Table 20** Statistical table for the effect of exogenous L-arginine concentration on quantification of residual serum L-Orn.

| Source of Variation | P value | P value summary | Significant? |
|---|---|---|---|
| **Interaction** | <0.0001 | **** | Yes |
| **Time** | <0.0001 | **** | Yes |
| **L-Arg +/- and Control (Com)** | <0.0001 | **** | Yes |

**Table 21** Samples comparison for the effect of exogenous L-arginine concentration on nitrite production and quantification of residual serum L-Arg, L-Cit and L-Orn

| Samples comparison Tukey multiple comparison | Nitrite | | | Serum L-Arg | | | Serum L-Cit | | | Serum L-Orn | | |
|---|---|---|---|---|---|---|---|---|---|---|---|---|
| | T=0 | T=24 h | T=72 h | T=0 | T=24 h | T=72 h | T=0 | T=24 h | T=72 h | T=0 | T=24 h | T=72 h |
| **Cont. Com vs. untreated** | ns | * | ** | **** | **** | **** | **** | ns | ns | ** | ** | **** |
| **Cont. Com vs. 400 μM** | ns | ns | ns | ns | ns | ns | ns | ns | ns | ns | ns | * |
| **Cont. Com vs. 800 μM** | ns | *** | ns | ns | ns | **** | ns | ns | ns | ns | ns | ns |
| **Cont. Com vs. No L-Arg** | ns | ns | ns | ns | * | **** | ns | ns | ns | ns | ns | ns |

**Tables 17-21** Statistical tables for the effect of exogenous L-arginine concentration on nitrite production and quantification of residual serum L-Arg, L-Cit and L-Orn obtained from cultured BNL CL2 cells grown in the presence of 0, 400 or 800 μM L-arginine for T=0, 24 or 72 h. Tables summarise two-way ANOVA followed by a Tukey multiple comparison test using GraphPad Prism 9.4.1. The stars indicate the levels of significance; ns = P > 0.05, * = P ≤ 0.05, ** = P ≤ 0.01, *** = P ≤ 0.001 and **** = P ≤ 0.0001.

**Table 22** Statistical table for the effect of exogenous L-arginine with L-NAME or SNAP addition on nitrite production**.**

| Source of Variation | L-Arg + L-NAME | | | L-Arg + SNAP | | |
|---|---|---|---|---|---|---|
| | P value | P value summary | Significant? | P value | P value summary | Significant? |
| **Interaction** | <0.0001 | **** | Yes | <0.0001 | **** | Yes |
| **Samples, conditions and time points** | <0.0001 | **** | Yes | <0.0001 | **** | Yes |
| **L-Arg +/- L-NAME/SNAP** | <0.0001 | **** | Yes | <0.0001 | **** | Yes |

**Table 23** Statistical table for the effect of exogenous L-arginine with L-NAME or SNAP addition on quantification of residual serum L-Arg.

| **Source of Variation** | **L-Arg + L-NAME** | | | **L-Arg + SNAP** | | |
|---|---|---|---|---|---|---|
| | **P value** | **P value summary** | **Significant?** | **P value** | **P value summary** | **Significant?** |
| **Interaction** | 0.0005 | *** | Yes | 0.0015 | ** | Yes |
| **Samples, conditions and time points** | 0.1033 | ns | No | <0.0001 | **** | Yes |
| **L-Arg +/- L-NAME/SNAP** | <0.0001 | **** | Yes | <0.0001 | **** | Yes |

**Table 24** Statistical table for the effect of exogenous L-arginine with L-NAME or SNAP addition on quantification of residual serum L-Cit.

| **Source of Variation** | **L-Arg + L-NAME** | | | **L-Arg + SNAP** | | |
|---|---|---|---|---|---|---|
| | **P value** | **P value summary** | **Significant?** | **P value** | **P value summary** | **Significant?** |
| **Interaction** | 0.0338 | * | Yes | 0.9426 | ns | No |
| **Samples, conditions and time points** | 0.0172 | * | Yes | 0.0001 | *** | Yes |
| **L-Arg +/- L-NAME/SNAP** | 0.0026 | ** | Yes | <0.0001 | **** | Yes |

**Table 25** Statistical table for the effect of exogenous L-arginine with L-NAME or SNAP addition on quantification of residual serum L-Orn.

| **Source of Variation** | **L-Arg + L-NAME** | | | **L-Arg + SNAP** | | |
|---|---|---|---|---|---|---|
| | **P value** | **P value summary** | **Significant?** | **P value** | **P value summary** | **Significant?** |
| **Interaction** | 0.2626 | ns | No | 0.9403 | ns | No |
| **Samples, conditions and time points** | 0.0002 | *** | Yes | 0.0014 | ** | Yes |
| **L-Arg +/- L-NAME/SNAP** | 0.1975 | ns | No | 0.3062 | ns | No |

**Table 26** Samples comparison for the effect of exogenous L-arginine with L-NAME addition on nitrite production

| Samples comparison Bonferroni's multiple comparisons | L-Arg addition and L-Arg + L-NAME addition |
|---|---|
| | **Amount of Nitrite** |
| **Cont. Com - 24 h** | **** |
| **Cont. Com - 72 h** | ns |
| **400 µM - 24 h** | ns |
| **400 µM - 72 h** | **** |
| **800 µM - 24 h** | **** |
| **800 µM - 72 h** | *** |
| **No L-Arg - 24 h** | ns |
| **No L-Arg - 72 h** | **** |

**Table 27** Samples comparison for the effect of exogenous L-arginine with L-NAME addition on residual L-Arg, L-Cit and L-Orn quantitation

| Samples comparison Bonferroni's multiple comparisons | Amount of L-Arg | Amount of L-Cit | Amount of L-Cit |
|---|---|---|---|
| **400 µM - 24 h** | **** | ** | ns |
| **400 µM - 72 h** | **** | ns | ns |
| **800 µM - 24 h** | **** | ns | ns |
| **800 µM - 72 h** | **** | ns | ns |

**Table 28** Samples comparison for the effect of exogenous L-arginine with SNAP addition on nitrite production

| **Samples comparison Bonferroni's multiple comparisons** | **L-Arg addition and L-Arg + SNAP addition** |
|---|---|
| | **Amount of Nitrite** |
| **Cont. Com - 6 h** | **** |
| **Cont. Com - 24 h** | **** |
| **400 µM - 6 h** | **** |
| **400 µM - 24 h** | **** |
| **800 µM - 6 h** | **** |
| **800 µM - 24 h** | **** |
| **No L-Arg - 6 h** | **** |
| **No L-Arg - 24 h** | **** |

**Table 29** Samples comparison for the effect of exogenous L-arginine with SNAP addition on residual L-Arg, L-Cit and L-Orn quantitation

| **Samples comparison Bonferroni's multiple comparisons** | **Amount of L-Arg** | **Amount of L-Cit** | **Amount of L-Cit** |
|---|---|---|---|
| **400 µM - 6 h** | ns | ns | ns |
| **400 µM - 24 h** | **** | * | ns |
| **800 µM - 6 h** | *** | * | ns |
| **800 µM - 24 h** | ns | ns | ns |

**Tables 22-29** The effect of exogenous L-arginine concentration on nitrite production and quantification of residual serum L-Arg, L-Cit and L-Orn obtained from cultured BNL CL2 cells grown in the presence of 0, 400 or 800 µM L-arginine and the control complete DMEM media with addition of either L-NAME (4mM) or SNAP (100 µM) in BNL CL2 cells across the time points either 24 and 72 h or 6 and 24 h. Tables summarise two-way ANOVA followed by a Bonferroni's multiple comparisons test using GraphPad Prism 9.4.1. The stars indicate the levels of significance; ns = P > 0.05, * = P ≤ 0.05, ** = P ≤ 0.01, *** = P ≤ 0.001 and **** = P ≤ 0.0001.
