## Supplementary Tables for "L-Arginine supplementation modulates L-Arg/NO metabolic processes and AMPK/ACC-1 signalling in BNL CL2 hepatocytes"

### Supplementary Materials

**Table S1** Primers used for real-time qPCR analysis of the target genes.

| Target | Gene | NCBI ID | Sequence (5'-3') |
|---|---|---|---|
| **ACC-1** | *Acaca* | NM_133360.3 | Forward: GGGTCAAGTCCTTCCTGCTC |
| | | | Reverse: TTCCACACACGAGCCATTCA |
| **AMPK** | *Prkaa1* | NM_001013367.3 | Forward: GGATCCATCAGCAACTATCG |
| | | | Reverse: TCGACTCCTCCCCTGTCGAC |
| **β-actin** | *Actb* | NM_007393.5 | Forward: AGCTGAGAGGGAAATTGTGCG |
| | | | Reverse: GCAACGGAAACGCTCATT |

**Table S2** List of antibodies used and sourced from different companies for Western blotting.

| Target protein | Type | Dilution | Source company | Catalog No |
|---|---|---|---|---|
| **Anti-ACC-1** | Primary | 1:1000 | GeneTex | GTX132081 |
| **Anti-ACC-1-P (Ser79)** | Primary | 1:1000 | GeneTex | GTX133974 |
| **Anti-AMPK** | Primary | 1:1000 | Cell Signaling | 2532 |
| **Anti-AMPK-P (Thr172)** | Primary | 1:500 | Cell Signaling | 2535 |
| **Anti-β-actin** | Primary | 1:1000 | Sigma | A5441 |
| **Anti-mouse IgG-Peroxidase** | Secondary | 1:5000 | Sigma | A4416 |
| **Anti-rabbit IgG-Peroxidase** | Secondary | 1:5000 | Sigma | A6154 |

**Table S3** HPLC gradient program for separation of L-Arg, L-Cit and L-Orn at flow rate 1.1 mL/min.

| **Mobile Phase (%)** | **Time (min)** | | | | | | | | | | |
|---|---|---|---|---|---|---|---|---|---|---|---|
| | 0 | 15 | 20 | 24 | 26 | 34 | 38 | 40 | 42 | 42.1 | 49 |

| | | | | | | | | | | | |
|---|---|---|---|---|---|---|---|---|---|---|---|
| **A** | 86 | 86 | 70 | 65 | 53 | 50 | 30 | 0 | 0 | 86 | 86 |
| **B** | 14 | 14 | 30 | 35 | 47 | 50 | 70 | 100 | 100 | 14 | 14 |
